# Isolation, Identification and Determination of Antibiogram Characteristics of *Escherichia Coli* from Calves in West Shewa Zone, Oromia, Ethiopia

**DOI:** 10.1101/2025.10.02.677497

**Authors:** Lalisa Chali Deressa

## Abstract

A cross-sectional study, conducted from December 2022 to April 2024 in West Shewa, Oromia, Ethiopia aimed to isolate and identify *Escherichia coli* from calves and determine its antimicrobial susceptibility patterns. Diarrheic calves were purposively sampled, while non-diarrheic calves were randomly sampled. *Escherichia coli* was isolated using standard bacteriological methods. Stx1 and eaeA virulence genes were detected by PCR. *Escherichia coli* susceptibility to 11 antibiotics was assessed via disk diffusion. The study data was analyzed using SPSS version 20. A total of 162 rectal fecal samples were collected 84 from diarrheic calves and 78 from non-diarrheic calves aged 0-4 months. *Escherichia coli* was isolated in 48.1% (78/162; 95% CI: 40.2–56.1%). *Escherichia coli* was more common in diarrheic calves (56%) than non-diarrheic ones (40%) (p = 0.039). Season, age, health, house hygiene, first colostrum feeding, and feeding technique all significantly impact on the occurrence of *Escherichia coli* (p < 0.05). 10 of the 78 isolates were PCR-tested for eaeA and stx1 genes. The PCR results showed 30% had eaeA, 60% had stx1, and 10% had both genes. Isolates from diarrheic (n=24) and non-diarrheic (n=16) calves showed resistance to ampicillin, amoxicillin, tetracycline, and amoxicillin/clavulanic acid, but were susceptible to gentamicin, norfloxacin, nitrofurantoin, and streptomycin. The 54.2% of isolates from diarrheic calves and 50% from non-diarrheic calves were multidrug resistant. It was concluded that *Escherichia coli* may cause calf diarrhea, influenced by season, age, health, and colostrum feeding timing. The effective antibiotics identified were gentamicin, nitrofurantoin, norfloxacin, and streptomycin. Further characterization of *Escherichia coli* in calves is needed to improve calf health and antibiotic stewardship.

## 1. INTRODUCTION

Calves are valuable resources for dairy and beef herd sustainability that they are a source of animal production used for breeding or meat [1]. However, they are suffering from higher mortality than their adult counterparts [2,3]. Diarrhea is a major disease that can be fatal to newborn calves and has multifactorial causes including *Escherichia coli* (*E. coli)*, *Salmonella*, rotaviruses, coronaviruses, and *Cryptosporidium* species [4]. *E. coli* is the major organism in the normal intestinal microbiota of mammals; that, it plays a vital role in host metabolism, immunology, and nutrition. However, sometimes it can cause mild-to-severe intestinal infections in humans and animals [5,6]. The enterotoxigenic *E. coli* (ETEC) is among the intestinal pathogenic *E. coli* (IPEC) to cause colibacillosis in young calves. However, enteropathogenic *E. coli* (EPEC), enteroinvasive *E. coli* (EIEC), and enterohemorrhagic *E. coli* (EHEC) have also been stated as causative agents of calf fatal coli septicemia [7].

Calf diarrhea caused by *E. coli* can lead to elevated morbidity and mortality rates, which resulted in large global economic losses. Besides calf death, other costs, such as treatment, diagnostics, prevention and control, and labor intervention costs, reduced herd size and chronic illnesses, which might result in impaired performance, contribute to economic losses to the [8].

Furthermore, cattle originated *E. coli* is a vital public health risks because cattle are a major reservoir of EHEC, EPEC, and Shiga toxin-producing *E. coli* (STEC), which are pathogens linked to food-borne diseases [9]. Humans can contract an *E. coli* infection if they consume something that has been contaminated with animal feces or come into contact with it. From a zoonotic perspective, STEC is the only group of pathogenic *E. coli* of significant interest, that the Shiga toxin-producing strains are able to cause severe disease in humans when being transmitted from the reservoir animals via the food chain [10,11]. The main source of *E. coli* O157:H7 is healthy cattle, which harbor this pathogen without showing any symptoms. Hence, *E. coli* in food animals has the potential to contribute to zoonotic infections [12].

The cause of calf diarrhea is multifactorial, with the main risk factors being poor calves management, age, breed, and feeding. *E. coli* has been isolated from calves that had diarrhea, and the risk factors associated to the isolates are identified as younger age and poor colostrum feeding [13,14]. Although, *E. coli* is susceptible to some antimicrobials, it is able to acquire and donate resistance genes through horizontal gene transfer [4]. These antimicrobial-resistant *E. coli* strains are mostly found in the gut of calves but they are usually not harmful to the calf itself. However, it can inflict life-threatening infections on other members of the herd [15]. The spread of *E. coli* antibiotic resistance genes from animals to people through food, direct contact, or shared environmental resources is a serious public health issue [16].

Calf diarrhea, particularly caused by *E. coli*, is a major problem in the livestock industry and can cause farmers to suffer large financial losses in Ethiopia [8,17]. Certain studies conducted in Ethiopia indicated that the prevalence of calf diarrhea caused by *E. coli* was 70.66 [18], 72% [19], and 80.77% [20]. As study reported by [7], severe diarrhea in calves caused by *E. coli* resulted calf fatality and poses a financial risk to the Ethiopia dairy production. The study also revealed that calf diarrhea has been reported at a prevalence of 15-20% and mortality rates of 1.5-8%. According to report by [17], diarrhea is a prevalent condition in calves younger than three months and is still a major cause of lost productivity and financial loss for cattle farmers. And also, it causes global increases in morbidity and mortality rates. It has consequently had a significant impact on the dairy industry globally. Moreover, an increase in the resistance of *E. coli* to commonly used antimicrobials has also been noted in both public health and veterinary sectors in Ethiopia [21, 22, 23]. The data from the European Antimicrobial Resistance Surveillance Network (EARS-Net) of 2019 showed that 44.2% of *E. coli* cases had antimicrobial resistance (AMR), and more than half of the *E. coli* isolates were resistant to at least one antimicrobial group [24]. Commensal *E. coli* in the gastrointestinal tract (GIT) of most healthy animals can act as reservoirs for horizontal transferable resistance genes and are used for the occurrence of resistance in *E. coli* [15]. The use of antibiotics in livestock, whether for growth promotion or disease treatment, contributes to the emergence of resistant bacteria [25]. This can complicate treatment regimens in animals which often incurs higher healthcare costs due to more expensive medications, and additional tests and procedures. These resistant strains can affect to human by transmitted from animals to humans through direct contact, ingestion of tainted animal products, or exposure to the environment [5].

Many studies conducted in Ethiopia have shown the importance of *E. coli* infection in diarrheic calves [14, 18, 6, 26], but few have looked at the incidence of *E. coli* infections, strains of pathogenic *E. coli* and its resistance pattern in diarrheic and non-diarrheic calves [7, 27]. Moreover, there is a dearth of studies conducted on the prevalence, pathotype of *E. coli* and its antimicrobial susceptibility test in both diarrheic and non-diarrheic calves within the West Shewa zone of the Oromia regional state in general and in Ambo, Bako Tibbe, and Ejere districts in particular. Therefore, the goals of this study were to isolate and identify pathogenic *E. coli* from diarrheic and non-diarrheic calves, ascertain the AMR pattern of the isolates and pinpoint the risk factors associated to *E. coli* infection, ensuring a full understanding of the problem in order to plan and achieve control strategies.

## 3. Material and Methods

### 3.1 Description of Study Area

The study took place from December 2022 to April 2024 in Ambo, Bakko Tibbe, and Ejere Districts of the West Shewa Zone. Ambo’s population is about 112,129, with 50.08% female and 49.92% male. Primarily practices mixed crop-livestock farming, housing 145,371 cattle, 50,152 sheep, 27,026 goats, 105,794 chickens, 9088 horses, 2914 donkeys, and 256 mules [28].

Bako Tibe district has a population of 133,799, comprising 68,401 males and 65,398 females, primarily engaged in mixed farming. The district’s livestock includes 137,343 cattle, 12,502 sheep, 15,381 goats, 96,742 chickens, 3,685 horses, 1,023 mules, and 8,415 donkeys [29].

Ejere district has an estimated population of 13,423, with 6,420 males and 7,003 females, primarily practicing mixed farming. The district’s livestock includes 91,770 cattle, 44,756 sheep, 9,127 goats, 18,787 donkeys, 360 mules, and 107,780 poultry [30].

**Figure 1:**
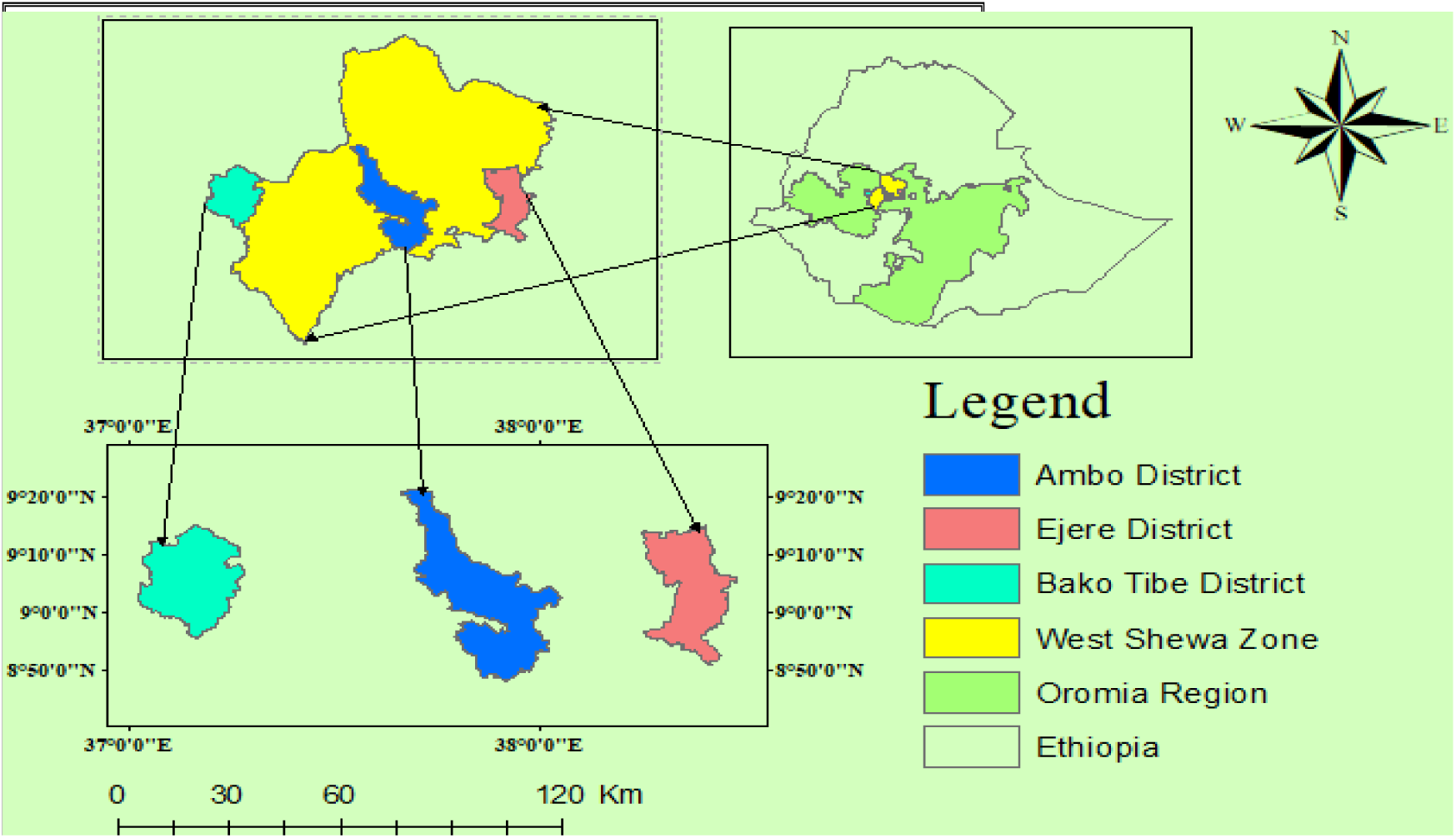
Map of Study Area

### 3.2 Study Population

The study populations comprised both diarrheic and non-diarrheic calves found in West Shewa Zone Ambo, Bakko Tibbe, and Ejere Districts. Cross-breed and local breed calves raised in an extensive, semi-intensive, and intensive farming system were taken as the target population. The study animals comprised calves of both sexes that were between 0 and 4 months old, according [8, 14, 20]. The ages of the calves were categorized into four age groups: less than 30 days, 30– 60 days, 61–90 days, and 91–120 days [31]. The calves’ body condition was recorded as good, medium, or poor [32]. Calves’ health status was recorded as diarrheic and non-diarrheic depending on whether or not they had diarrhea [33]. Also, there were the two-time category of first colostrum feeding: those who received it before and after three hours [34].

### 3.2 Study Design and Sampling Method

A cross-sectional study was conducted on a total of 162 calves, of which 84 were diarrheic, and the remaining 78 were non-diarrheic. The diarrheic calves were sampled purposively based on the availability of cases of diarrhea in calves and the willingness of the farm owners; whereas the non-diarrheic calves were randomly selected. Every calf underwent a physical examination to assess its health, and the calves were assigned as diarrheic if they had paste-like consistency feces, and their tails and perineum were dirty with feces. Rectal fecal samples were collected after digital stimulation of the rectal mucosa using a one-use rubber glove.

### 3.3 Sample Size Determination

The sample size for non-diarrheic calves was based on a 4.6% expected prevalence from previous reports at Alage Dairy Farm, Ethiopia [7].

N = p (1-p) (z/E) ^2^, Where: n = required sample size; p = expected prevalence; and Z = the zero score (95% CI = 1.96) and E = the margin of error (at 95% CI = 0.05).

This formula indicated that 67 non-diarrheic calves were intended to be included in the study. However, in order to increase the accuracy of a prevalence estimate in non-diarrheic calves, the sample size was increased by 15%, and a total of 78 non-diarrheic calves were added to the total number of 84 diarrheic calves. Therefore, 168 calves in all were sampled for the study.

### 3.4 Questionnaire Survey

Farm management systems were evaluated through a structured questionnaire distributed to dairy farm owners to address the study’s objectives. The text addresses key aspects of calf husbandry, including health care, farm management, hygiene, and colostrum feeding practices. Farm owners were interviewed in person in Afan Oromo and Amharic (Annex 1).

### 3.5 Sample Collection and Processing

A sufficient amount (25–50 g) of fecal samples was collected directly from the rectum of each calf by manual stimulation using disposable latex gloves. The sample was then put into a sterile, 50 ml wide-mouth screw-capped, universal bottle. The samples were labeled with the date of collection, district, and sample ID and transported to Ambo University Zoonosis and Food Safety Laboratory under ice packs in an ice box for *E. coli* isolation. The samples were processed as soon as possible upon arrival at the laboratory and, if not stored in refrigeration at +4 °C [35]. Upon sampling, the samples were documented in a data recording format with the following details: the calves’ age, breed, sex, origin, sampling date, etc. (Annex 2).

### 3.6 Isolation of the *E. coli*

#### 3.6.1 Bacteriological Culturing and Examination

Fecal samples enriched in peptone water were incubated at 37℃ for 24 hours, then cultured on MacConkey agar. Pink colonies, indicating lactose fermentation, were suspected to be *E. coli*. Suspected *E. coli* colonies were sub-cultured on Eosin Methylene Blue (EMB) agar and incubated at 37℃ for 18–24 hours. Colonies exhibiting a green metallic sheen were identified as *E. coli* and transferred to nutrient agar for further biochemical testing after 24 hours at 37℃ [13]. The Laboratory test results were documented in the recording format (Annex 3).

#### 3.7.2. Biochemical Tests

The representative colonies were sub-cultured on the nutrient agar and incubated at 37℃ for 24 hours until pure colonies were obtained for further biochemical tests. The biochemical tests like Triple Sugar Iron (TSI) and IMViC were conducted according to the protocols described under Annex 5. Briefly, the TSI test was carried out in such a way that the *E. coli* suspected colony was inoculated into TSI slant agar. After incubating the inoculated media at 37°C for 24 hours, the isolates that produced a yellow slant and butt with gas production but no hydrogen sulfide production were taken to be *E. coli*. The IMVic test was used to biochemically confirm *E. coli* using indole and methyl red positivity and Voges-Proskauer and citrate negativity [36] (Annex 10, Figure 7). The biochemically confirmed *E. coli* isolates were grown in nutrient broth and mixed with an equal volume of 20% glycerol, and preserved in the refrigerator at -20℃ until molecular and drug susceptibility tests carried out.

#### 3.7.3. Molecular Detections in the National Veterinary Institute (NVI)

Ten selected *E. coli* (7 from diarrheic and 3 from non-diarrheic) isolates were tested for virulence genes using multiplex PCR [37]. A multiplex PCR assay was employed to examine virulence genes and their corresponding pathotypes in diarrheagenic *E. coli*. Virulence genes, such as eaeA for EPEC pathotype, eaeA with stx (1 or 2) for EHEC pathotype, and Stx1 or Stx2 for STEC pathotype were assessed [38]. The base sequences, primer name, target gene, and anticipated amplified product sizes for each unique oligonucleotide primer used in the study (Table 1).

**Table 1:**
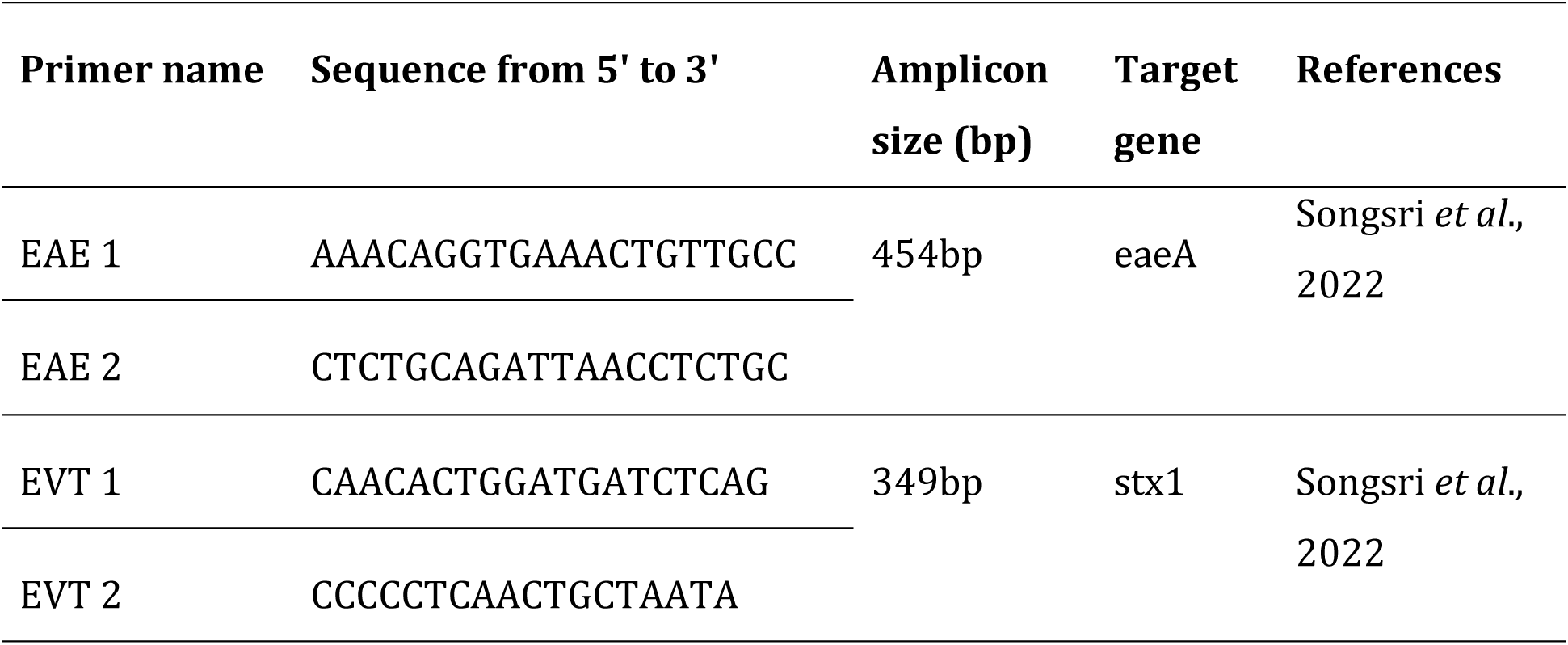
Primers, Sequences, and Amplicon Size of *E. coli* genes.

##### 3.7.2.1 DNA extraction

The DNAs from *E. coli* isolates were extracted using the boiling method. In brief, 200μL of overnight broth culture of each isolate was put into a 1.5 ml microcentrifuge tube, and 200μL buffer solution of AL and 20μL of protease were added. The mixture was then pulse-vortexed for 15 seconds and incubated at 56 °C for 10 minutes. It has been expected that DNA yield will reach a maximum after lysis for 10minutes at 56°C. Lastly, 200µl of ethanol (96–100%) was added to the mixture, and mixed by pulse-vertexing. The mixture in 1.5 ml microcentrifuge tube was then quickly centrifuged at 6000xg (8000rpm) for 1 min. The suspension was carefully transferred to the QIA amp spin column (a 2 ml collection tube) without wetting the rim; the cap was closed and centrifuged at 6000 x g (8000 rpm) for 1 minute. The supernatant in the collection tube was discarded and the tube was then in a clean 2 ml collection tube. The tube was then boiled and immediately placed on ice and centrifuged at 14,000 rpm for 3 minutes. In order to use the supernatant as template DNA, it was collected and kept at -20 °C. Ultraviolet absorbance spectrometry used to measure the quality of DNA.

##### 3.7.3.2 Gene amplification

For PCR amplification, the thermal cycling protocol comprised 5 minutes of pre-denaturing at 94°C for eaeA and 95°C for stx-1 gene, 35 cycles of denaturing for one minute at 94°C for eaeA and 95°C for stx-1, annealing for 90 seconds at 55°C, extending for one minute at 72°C, and concluding with a final extension for 7 minutes at 72°C for eaeA and stx-1 genes.

##### 3.7.3.3 Electrophoresis

The PCR products were then examined using the agarose gel electrophoresis technique. Then stained with a 2% agarose made in 4 μl Gel red with loading dye using a 10 µl markers DNA ladder of 349 (stx-1) and 454 (eaeA) base pairs (bp) 10 µl markers (ladder) and 10µl PCR products, then allowing to run it to 120 volts for 60 minutes. The samples were also placed with known positive and negative controls for the shigatoxin-1 (stx-1) and intimin (eaeA) genes. Finally, the PCR products were viewed using a UV transilluminator.

### 3.8. Antimicrobial Susceptibility Test

The antimicrobial susceptibility testing of *E. coli* isolates was conducted using Kirby Bauer’s disc diffusion method on Mueller Hinton agar [40]. According to [19], a turbidity standard of 0.5 McFarland was used to estimate the density of a bacterial suspension. The antimicrobials tested were amoxicillin (2μg), amoxicillin/clavulanic acid (10μg), ampicillin (10μg), ciprofloxacin (5μg), gentamicin (10μg), nalidixic acid (30μg), nitrofurantoin (300μg), norfloxacin (10μg), streptomycin (10μg), tetracycline (30μg) and trimethoprim (5μg). These antimicrobials were chosen due to their efficacy and accessibility in treating *E. coli* infections in humans and animals. The antibiotic inhibition zone diameters (IZD) were measured in millimeters using a caliper. The IZD were compared with the standard given by [40] and interpreted as susceptible, intermediate, or resistant. The isolates designated as MDR were demonstrated based on the ability of the tested isolates to resist the effects of three or more antibiotics. Antibiotic susceptibility test protocols stated in Annex 8.

### 3.9 Data Management and Analysis

All biological and survey data were entered into a Microsoft Excel spreadsheet and statistically analyzed using SPSS software version 20. Descriptive statistics such as frequencies and percentages were used to analyze, and the results were displayed in tables. The associations between the prevalence of *E. coli* and the risk factors were examined using the Pearson’s χ2 test. In all of the analysis, the 95% confidence interval with a significance of p-value <0.05 was set to indicate the presence of significance.

## 4. Results and Discussion

### 4.1. Results

#### 4.1.1. Prevalence of E. coli in Calves

Out of 162 calf fecal samples (84 diarrheic, 78 non-diarrheic), 78 (48.1%) tested positive for *E. coli* (95% CI: 40.2–56.1%). *E. coli* prevalence was 56% in diarrheic calves and 39.7% in non-diarrheic calves, with a significant difference between groups (p < 0.05) (Table 2).

**Table 2:** Overall Prevalence of *E. coli* in Diarrheic and Non-diarrheic calves.

| Healthy status | No.<br>examined | No<br>positive | Prevalence % | Pearson's $\chi^2$ and p-value |
| --- | --- | --- | --- | --- |
| Diarrheic calves | 84 | 47 | 56.1 | 0.039 |
| Non-diarrheic<br>calves | 78 | 31 | 39.70 |  |
| **Total** | **162** | **78** | **48.10** |  |

#### 4.1.2. Risk Factor associated with E. coli prevalence in calves

The expected risk factors associated with the prevalence of *E. coli* were breed, sex, and origin of the animal, seasons (Autumn (April - May), Summer (June – August) and Spring (September - October)), health status of the calves, body condition, farm husbandry, farm size, house hygiene, colostrum feeding time, and colostrum feeding methods. Accordingly, the analysis using Pearson Chi-square (χ^2^) test revealed that the isolation rate of *E. coli* was significantly correlated with age, health status, colostrum feeding times, colostrum feeding methods and origin of the calves, and seasons, and, house hygiene, (P≤0.05). On the contrary, no significant association was found between breed, sex, body condition, farm husbandry, and farm size (P > 0.05) (Table 3).

**Table 3:** Pearson’s χ2 results on *E. coli* positivity and related risk factors.

| Variable<br>s | Category | No. examined | Positive sample % | $\chi^2$ | *P-value* |
| --- | --- | --- | --- | --- | --- |
| Season | Autumn | 43 | 19(44.18) | 11.843 | 0.003 |
|  | Summer | 77 | 47(61.00) |  |  |
|  | Spring | 42 | 12(28.57) |  |  |
| Sex | Male | 54 | 22(40.74) | 1.780 | 0.182 |
|  | Female | 108 | 56(51.90) |  |  |
| Age in<br>days | < 30 | 14 | 10(71.40) | 11.283 | 0.010 |
|  | 30 – 60 | 20 | 15(75.00) |  |  |
|  | 61 – 90 | 41 | 16(39.00) |  |  |
|  | 91 -120 | 87 | 37(42.50) |  |  |
| Breed | Cross breed | 149 | 71(47.70) | 0.184 | 0.668 |
|  | Local | 13 | 7(53.80) |  |  |
| Body<br>condition<br>s | Good | 38 | 17(44.70) | 3.637 | 0.162 |
|  | Medium | 83 | 36(43.40) |  |  |
|  | Poor | 41 | 25(61.00) |  |  |
| Districts | Bako Tibe | 38 | 14(36.80) | 6.474 | 0.039 |
|  | Ambo | 63 | 27(42.90) |  |  |
|  | Ejere | 61 | 37(60.70) |  |  |
| Healthy<br>status | Diarrheic | 84 | 47(56.00) | 4.256 | 0.039 |
|  | Non-diarrheic | 78 | 31(39.70) |  |  |
| Farm<br>husbandr<br>y | Intensive | 118 | 57(48.30) | 0.277 | 0.870 |
|  | Semi-intensive | 33 | 15(45.50) |  |  |
|  | Extensive | 11 | 6(54.50) |  |  |
| Farm size | Large | 39 | 18(46.20) | 0.138 | 0.934 |
|  | Medium | 89 | 44(49.40) |  |  |
|  | Small | 34 | 16(47.10) |  |  |
| House<br>hygiene | Good | 24 | 12(50.00) | 7.425 | 0.024 |
|  | Medium | 93 | 37(39.80) |  |  |
|  | Poor | 45 | 29(64.40) |  |  |
| CFT | < 3 hours | 130 | 53(40.80) | 14.353 | 0.001 |
|  | > 3 hours | 32 | 25(78.10) |  |  |
| MCF | Suckling | 122 | 51(41.80) | 7.967 | 0.005 |
|  | Hand feeding | 40 | 27(67.50) |  |  |

##### 4.1.2.1 Virulence genes and strains of *E. coli*

Ten of the 78 *E. coli* isolates were tested for virulence genes (stx1 and eaeA) via PCR. All 10 (100%) contained at least one gene, and 1 (10%) had both genes (Table 4).

**Table 4:** PCR detected *E. coli* virulence genes.

| Virulent gene detected | No. of isolates Examined | No. of genes examined | % of genes detected |
| --- | --- | --- | --- |
| eaeA | 10 | 3 | 30 |
| stx-1 | 10 | 6 | 60 |
| eaeA and stx-1 | 10 | 1 | 10 |
| Total | 10 | 10 | 100 |

The virulence genes were detected in both isolates obtained from diarrheic and non-diarrheic calves. Of the 7 isolates from diarrheic calves tested for the genes, 3 (42.86%) carried eaeA, 3 (42.86%) carried stx1, and 1 (14.29%) carried both eaeA and stx1 virulence factors. In the case of the three isolates from non-diarrheic calves tested for the genes, all (100%) tested contained only stx1, but none of them harbored eaeA (Table 5).

**Table 5:**
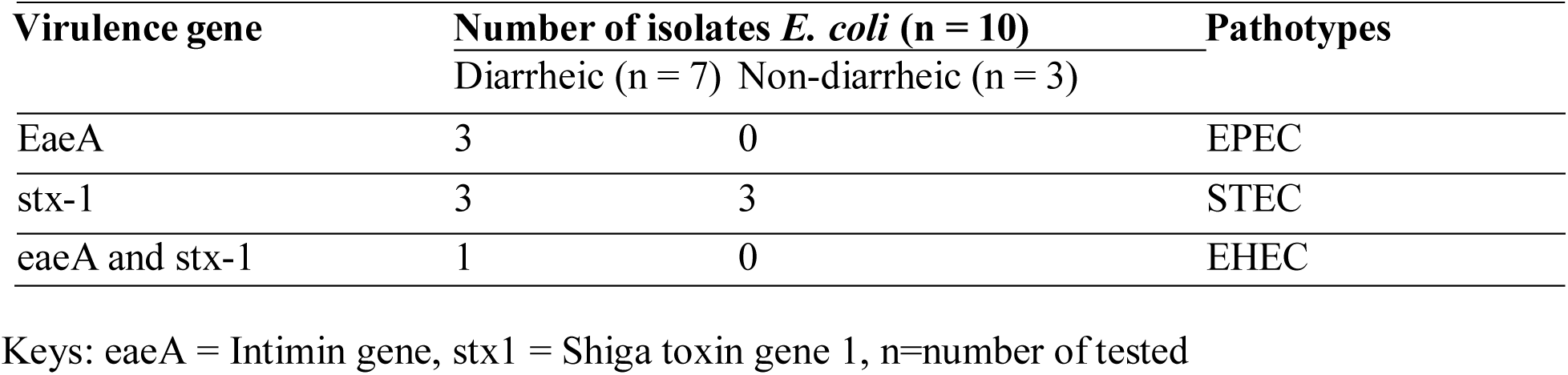
Virulence genes detected and health status of the calves.

| Virulence gene | Number of isolates *E. coli* (n = 10) |  | Pathotypes |
| --- | --- | --- | --- |
|  | Diarrheic (n = 7) | Non-diarrheic (n = 3) |  |
| EaeA | 3 | 0 | EPEC |
| stx-1 | 3 | 3 | STEC |
| eaeA and stx-1 | 1 | 0 | EHEC |

The virulence genes discovered were used to identify the pathotypes of *E. coli* depending on their virulence genes detected. The *E. coli* isolates that were tested positive for stx1 without the presence of eaeA are referred to as STEC; whereas the isolates that were tested positive for eaeA were EPEC, or those who tested positive for both stx1 and eaeA were EHEC (Table 6).

**Table 6:** *E. coli* Pathotypes from diarrheic and non-diarrheic calves.

| Virulence gene | Number of isolates *E. coli* (n = 10) |  | Pathotypes |
| --- | --- | --- | --- |
|  | Diarrheic (n = 7) | Non-diarrheic (n = 3) |  |
| eaeA | 3 | 0 | EPEC |
| stx-1 | 3 | 3 | STEC |
| eaeA and stx-1 | 1 | 0 | EHEC |

**Figure 2:**
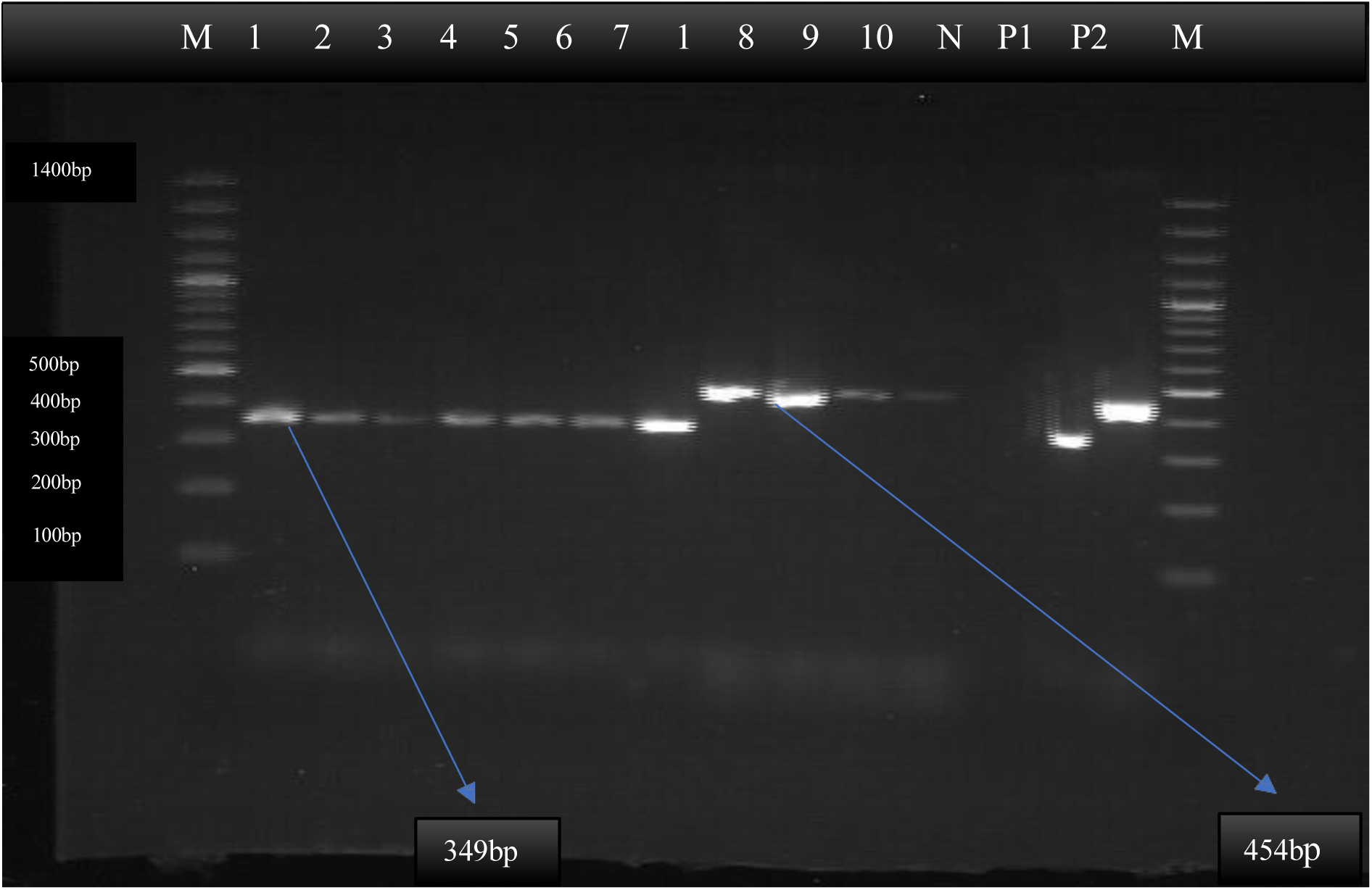
Agarose gel electrophoresis of amplified eaeA and stx1 gene 2% Agarose gel electrophoresis of amplified eaeA and stx1 genes generating a 454 and 349 base pair amplicon, respectively. M = Molecular ladder started 100bp, N = Negative control, P1 = Positive control for shigatoxin-1 (*stx1*) gene, P2 = Positive control for intimin (*eaeA)* gene, Lane no 1 2 3 4 5 6 7 8 9 and 10 indicates sample no. Lanes 1 - 7 are positive results for stx1, and 1, 8, 9, and 10 are positive results for intimin.

##### 4.1.2.2 Antimicrobial susceptibility test

Of the 78 *E. coli* isolates, 40 (24 from diarrheic and 16 from non-diarrheic calves) were tested for antimicrobial susceptibility against 11 commercially available antimicrobial disks. The results of the test showed that the majority of *E. coli* isolates from diarrheic and non-diarrheic calves were resistant to ampicillin (10μg) 100% in diarrheic and non-diarrheic calves, Amoxicillin (2μg) 91.7% in diarrheic and 100% in non-diarrheic calves, Tetracycline (30μg) 87.5% in diarrheic and 93.75% non-diarrheic calves, and amoxicillin/clavulanic acid (10μg) 79.17% in diarrheic and 87.5% in non-diarrheic calves. Conversely, the isolated from both diarrheic and in non-diarrheic calves were highly susceptible to gentamicin (10μg) 91.7% in diarrheic and 100% non-diarrheic calves, norfloxacin (10μg) 83.3% in diarrheic and 93.75% in non-diarrheic calves, and nitrofurantoin (300μg) 79.17% in diarrheic and 93.75% in non-diarrheic calves (Table 7).

**Table 7:** AMR of *E. coli* in diarrheic and non-diarrheic calves.

| Antimicrobials and<br>Concentration µg/disc | Antimicrobial<br>classes | Number of *E. coli* Isolates (n = 40) |  |  |  |  |  |
| --- | --- | --- | --- | --- | --- | --- | --- |
|  |  | Diarrheic Calves (n= 24) |  |  | Non-diarrheic calves (n = 16) |  |  |
|  |  | S (%) | I (%) | R (%) | S (%) | I (%) | R (%) |
| Amoxicillin (2 µg) | Beta-lactams | 1(4.17) | 1 (4.17) | 22 (91.7) | 0(0) | 0(0) | 16 (100) |
| Amoxicillin/clavulanic acid (10 µg) | Beta-lactams | 1 (4.17) | 4(16.67) | 19(79.17) | 0(0) | 2(12.5) | 14(87.5) |
| Ampicillin (10 µg) | Beta-lactams | 0(0) | 0(0) | 24 (100) | 0(0) | 0(0) | 16 (100) |
| Ciprofloxacin (5µg) | Quinolone | 7(29.17) | 8 (33.3) | 9 (37.5) | 5(31.25) | 5(31.25) | 6 (37.5) |
| Nalidixic acid (30µg) | Quinolone | 10(41.67) | 11(45.83) | 3 (12.5) | 1(6.25) | 14(87.5) | 1(6.25) |
| Norfloxacin (10µg) | Quinolone | 20 (83.3) | 3 (12.5) | 1 (4.17) | 15(93.75) | 1 (6.25) | 0(0) |
| Nitrofurantoin (300µg) | Nitrofuran | 19(79.17) | 2 (8.33) | 3 (12.5) | 15(93.75) | 1(6.25) | 0(0) |
| Gentamicin (10µg) | Aminoglycosides | 22 (91.7) | 1 (4.17) | 1 (4.17) | 16 (100) | 0(0) | 0(0) |
| Streptomycin (10µg) | Aminoglycosides | 17 (70.83) | 5(20.83) | 2 (8.33) | 9 (56.25) | 3(18.75) | 4 (25) |
| Tetracycline (30µg) | Tetracyclines | 0(0) | 3 (12.5) | 21(87.5) | 0(0) | 1(6.25) | 15(93.75) |
| Trimethoprim (5µg) | Folate pathway<br>inhibitors | 14 (58.33) | 8(33.33) | 2 (8.33) | 7(43.75) | 7(43.75) | 2(12.5) |

The multidrug resistance pattern of the *E. coli* isolates disclosed that of the 24 isolates from diarrheic calves, 13 (54.2%) showed resistance to at least three antimicrobial agents. The highest percentage of MDR was to three drugs (37.5%) Table 8.

**Table 8:** Multidrug resistance *E. coli* isolates from Diarrheic calves.

| No of antibiotics | AMR pattern | No. of isolates | Resistant (%) |
| --- | --- | --- | --- |
| One | AMP | 2 | 8.33 |
| Two | AMP, TET | 9 | 37.5 |
| Three | AMP, S, TMP (1) | 9 | 37.5 |
|  | AMP, TET, TMP (1) |  |  |
|  | AMP, CIP, TET (4) |  |  |
|  | AMP, S, TET (1) |  |  |
|  | AMP, NAL, TET (1) |  |  |
|  | AMP, NOR, TET (1) |  |  |
| Four | AMP, CIP, NIT, TET (3) | 4 | 16.67 |
|  | AMP, CIP, GEN, TET (1) |  |  |

Keys: AMP=Ampicillin, TET = Tetracycline, CIP = Ciprofloxacin, TMP = Trimethoprim, S = Streptomycin, NAL=Nalidixic acid, NIT=Nitrofurantoin, GEN=Gentamycin, NOR=Norfloxacin

Regarding the *E. coli* isolates from non-diarrheic calves, of the 16 isolates tested, 8 isolates (50%) were resistant to at least three antimicrobial agents. The highest percentage of multidrug resistance was to three drugs (37.5%) (Table 9).

**Table 9:**
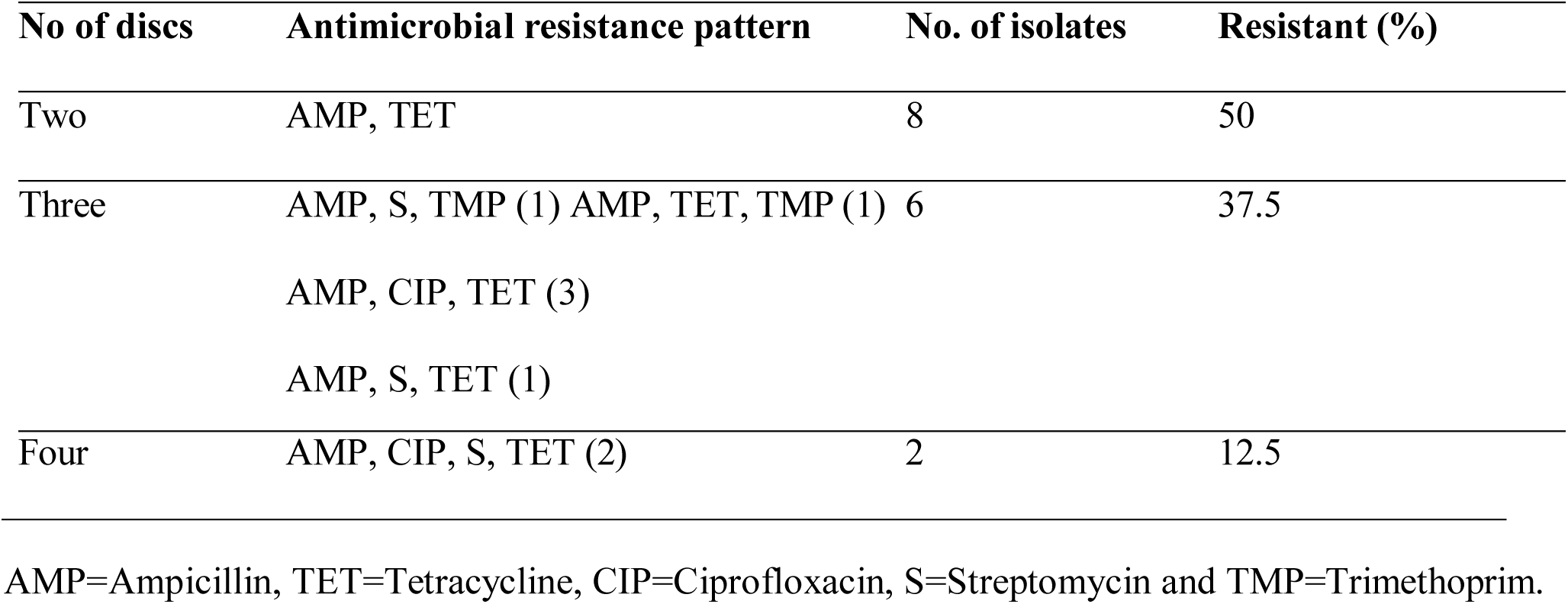
Multidrug resistance of *E. coli* from non-diarrheic calves.

| No of discs | Antimicrobial resistance pattern | No. of isolates | Resistant (%) |
| --- | --- | --- | --- |
| Two | AMP, TET | 8 | 50 |
| Three | AMP, S, TMP (1) AMP, TET, TMP (1) | 6 | 37.5 |
|  | AMP, CIP, TET (3) |  |  |
|  | AMP, S, TET (1) |  |  |
| Four | AMP, CIP, S, TET (2) | 2 | 12.5 |

AMP=Ampicillin, TET=Tetracycline, CIP=Ciprofloxacin, S=Streptomycin and TMP=Trimethoprim.

### 4.1 Discussion

In the current study, the overall prevalence of *E. coli* in 0 to 4-month calves were 48.15%. The occurrences of the bacterium in diarrheic and non-diarrheic calves were 55.95% and 39.74%, respectively. Previous studies conducted by [13, 41] revealed a comparable result, with an overall prevalence of 46.4% in diarrheic and 49% in non-diarrheic calves. The overall prevalence in the current study was higher than the 8.11% previously reported from dairy calves in Ethiopia [7]. Compared to [42] at 56.3% and [43] at 64.5%, the current prevalence of 48.1% is lower. In India, *E. coli* the overall prevalence in calf feces was very high at 96.15%. In comparison, the percentages for diarrheic and non-diarrheic cases were 96% and 100%, respectively [44]. The reason why this study differs from previous reports could be attributed to differences in sample size and farm management practices. The small sample size of 52 rectal swabs from calves may limit insights into the true relationships between factors, especially given the varying farming practices among different farmers [15].

Many variables were examined in light of the risk factors linked to the isolation rate of *E. coli* in the current study, and the Pearson χ^2^ indicated that statistically significant relationships. The study revealed a significant seasonal difference (P<0.05) in *E. coli* isolation rates from calf diarrhea, with 28.6% in spring compared to 61% in summer. An Egyptian report found 66.6% of samples positive for *E. coli* in summer, similar to the 61% prevalence we observed during the rainy season [45]. As in the present study, the highest prevalence of *E. coli* (81.94%) was reported by [46] in the feces of calves during the rainy season in India. The [47] found that higher calving rates and cold stress, which increase calves’ susceptibility to *E. coli*, as well as the optimal humidity and temperature for bacterial growth, lead to the highest *E. coli* prevalence in summer.

The study found significant differences in *E. coli* presence across calf age groups (p = 0.010). Younger calves under 60 days had higher prevalence (71.4% under 30 days; 75% 30–60 days) compared to older calves (39% 61–90 days; 42.5% 91–120 days). In this study, 71.4% of calves were under 30 days old, surpassing the 20.8% reported in southern Ethiopia [7]. Our findings support [6] that younger calves are more susceptible to *E. coli* infections than older ones.

Calves infected with *E. coli*, whether exhibiting diarrhea or not, showed a significant association (p < 0.05). Consistent with the findings of [14], the higher prevalence of *E. coli* in diarrheic calves (56%) compared to healthy calves (39.7%) in this study highlights *E. coli* as a significant contributing factor to calf diarrhea. Our study found 56.0% of diarrheic calves infected with *E. coli*, aligning with [48] 51.8% in Jimma, Ethiopia. The higher occurrence of *E. coli* in diarrheic calves observed in this study is consistent with the findings of [49]. This increase is likely due to pathogenic strains such as ETEC, EPEC, EHEC, and STEC, which are known to cause diarrhea in calves. A study found a higher *E. coli* prevalence (33.33%) in diarrheic calves compared to 4.62% in healthy ones, which supports our findings of 56% in diarrheic and 39.70% in non-diarrheic calves [7]. This shows distinct *E. coli* strains cause intestinal and extraintestinal infections, with coli septicemic *E. coli* responsible for the latter [50]. The current study shows lower prevalence in diarrheic (56.0%) and healthy calves (39.7%) compared to reported by [43], who found 68.3% and 56.7%, respectively. Similarly, [42] reported a comparable prevalence in diarrheic calves (54.1%) but a higher rate in non-diarrheic calves (58.3%) than our findings. The differences in *E. coli* prevalence observed between this study and earlier research may be attributed to variations in management and hygiene practices across dairy farms [15]. Also, the use of random sampling reduces bias and enhances the accuracy of generalizing findings to the wider population.

Our results show that calves fed colostrum after three hours are significantly (p<0.05) more likely to develop *E. coli* infection than those fed within three hours, emphasizing the importance of early colostrum intake in preventing infection. A study discovered that calves fed within three hours of birth received a greater transfer of passive immunity than those fed after three hours [51]. According to [52], delayed and insufficient intake of high-quality colostrum hampers the transfer of essential passive immunity to the calf. Feeding calves colostrum immediately after birth ensures higher passive immunity transfer than feeding after 4 hours, due to declining antibody absorption after 6 hours [53]. This study confirms that calves receiving first colostrum within three hours of birth have lower early mortality, consistent with previous findings [34].

The polymerase chain reaction confirmed the presence of *E. coli* isolates with high sensitivity and specificity. The results also identified the distribution of key virulence genes among the strains: 30% were identified as EPEC, 60% as STEC, and 10% as EHEC. This study’s findings EPEC (30%), STEC (60%), and EHEC (10%) align with [27], who reported a 30% prevalence of EPEC in Ethiopia. This study reveals higher prevalence rates of STEC, EPEC, and EHEC compared to previous research. Specifically, the study reported lower prevalence rates in Ethiopia 13% for STEC, 6% for EPEC, and 6% for EHEC [23] while in India reported even lower rates: 6.1% for STEC, 2.9% for EPEC, and 0.3% for EHEC [4]. The study’s STEC prevalence (63.63%) aligns with report in Brazil [54], but is lower than the 100% EPEC reported by Hosein *et al*. (2019) and the 67.6% EHEC found in Egypt [45].

The virulence genes eaeA and stx1, which are associated with pathogenicity in *E. coli*, were detected in isolates from both diarrheic and non-diarrheic calves. While no strong link was observed between calf health and virulence genes, stx-1 was present in all healthy calves, whereas diarrheic calves also carried eaeA alongside stx-1. This aligns with studies which found a high prevalence of STEC in non-diarrheic calves [56, 4]. This is because bovines are the primary animal reservoirs for STEC, and most colonized animals do not show any symptoms while shedding the bacteria in their feces [57]. However, the current study did not corroborate the findings of [58], who reported that STEC was more frequently detected in diarrheic calves compared to non-diarrheic calves. This is because STEC strains may cause diarrhea in young calves due to inadequate colostrum intake and poor management [59].

Diarrheic calves harbored all three strains STEC, EHEC, and EPEC with STEC and EPEC most prevalent, unlike non-diarrheic calves, which only carried STEC. This study confirmed the presence of the Stx1 gene, indicating EHEC bacteria, and the intimin gene, indicating EPEC bacteria, in calves suffering from diarrhea. This aligns with the report identifying Stx1, Stx2, and intimin genes in diarrheic calves, confirming EHEC and EPEC as pathogenic types [60]. In contrast to [56], this study found that healthy calves are less likely to carry STEC, EHEC, and EPEC than diarrheic calves. This may be due to STEC, EHEC, and EPEC can be carried silently by healthy young calves, who are natural reservoirs for diarrheagenic *E. coli* strains [61].

Antimicrobial drug resistance in bacterial pathogens is an emerging concern. Our study found that over half of *E. coli* isolates from calves, whether diarrheic (54.2%) or non-diarrheic (50%), exhibited multidrug resistance. The high percentage of MDR (37.5%) in non-diarrheic calves may stem from *E. coli* in their normal gut flora, which can transfer antibiotic resistance genes [62]. Previous studies, such as [63], have also reported that antimicrobial-resistant *E. coli* can colonize calves as early as birth, aligning with the highest MDR observed in non-diarrheic calves in this study. The MDR prevalence in diarrheic calves in this study (54.2%) aligns with previous reports 49.6% in neonatal calves in India [4] and 56.7% in Ethiopian dairy farms [64]. The MDR in the current study (54.2%) is significantly lower than previous Ethiopian reports, which found prevalence of 90% and 84.6% among diarrheic newborns and children, respectively [48, 64]. Several Ethiopian studies report MDR rates at or near 100%, exceeding the current study. These include [66] in goats, [67] in animal-derived foods, in backyard chickens [68], in nursing cows and farm environments [26], in milk and dairy products [69], and in dairy cattle and humans [70]. The MDR in this study (54.2%) exceeds previous reports from Egypt (20.83%) [71] and Ethiopia (42.86%) by [72], indicating a higher resistance level. This study suggests that the high prevalence of MDR *E. coli* may result from widespread broad-spectrum antibiotic use, inappropriate prescribing, and bacterial genetic mutations [26, 70]. In this study, *E. coli* isolates from both healthy and diarrheal calves exhibited resistance to at least one antimicrobial tested. *E. coli* from diarrheic calves showed high resistance to ampicillin, amoxicillin, tetracycline, and amoxicillin/clavulanic acid, but remained sensitive to gentamicin, norfloxacin, nitrofurantoin, and streptomycin, with lower sensitivity to trimethoprim (Table 4.8). This study confirms previous findings by [71] that *E. coli* from diarrheic calves is resistant to tetracycline and amoxicillin but susceptible to gentamicin, streptomycin, and trimethoprim. It also aligns with [19] in showing sensitivity to streptomycin, though it contradicts their report of high tetracycline sensitivity, as our isolates exhibited high resistance. In this study, 87.5% of *E. coli* isolates were resistant to tetracycline, contrasting with [18], who found the bacteria highly sensitive to it. The current study found that 70.83% of *E. coli* isolates were susceptible to streptomycin, contrasting with [8] findings of 74% resistance. Both studies, however, reported high resistance to ampicillin and tetracycline.

All non-diarrheic calf *E. coli* isolates showed resistance to amoxicillin and ampicillin, with most also resistant to tetracycline and amoxicillin/clavulanic acid. They remained sensitive to gentamicin, nitrofurantoin, and norfloxacin, with only a few sensitive to streptomycin (Table 4.9). Comparable outcomes were observed in the earlier studies [26], wherein all isolates were determined to be resistant to ampicillin (100%) susceptibility against gentamicin (100%). The current findings concur with the earlier research conducted in China [74], that demonstrated the *E. coli* isolates from healthy cattle were extremely resistant to ampicillin, tetracycline, and amoxicillin/clavulanic acid. The present results were also in line with reports from Bangladesh [75], which exhibited the most resistance *E. coli* isolates against amoxicillin, ampicillin, and tetracycline from dairy milk. Similarly, the current study confirmed the findings of the studies regarding the high resistance of non-diarrheic *E. coli* isolates to tetracycline and ampicillin from calves [76, 63]. However, the earlier report, which showed 100% susceptibility against tetracycline, was not consistent with the current study, which found that all isolates were resistant to tetracycline. The present study, which corroborated previous findings by [44], discovered that isolates from healthy calves exhibited increased resistance to ampicillin and tetracycline, but gentamicin remained sensitive. Moreover, the variations in antibiotic sensitivity of *E. coli* in different areas could be due to numerous factors, including treatment selection, antibiotic overuse, and animal species differences, as well as the pathogen can spread resistance genes by horizontal gene transfer [18, 77]. This may also be connected to the fact that district veterinary clinics in Ethiopia treat their animal patients with common antimicrobials based on tentative disease diagnoses [68].

The information from the current study, which examines the isolation and identification of *E. coli* in diarrheic and non-diarrheic calves as well as determining its antimicrobial susceptibility, could be utilized for a number of purposes. It provides a clue regarding the identification and frequency of *E. coli* isolated from diarrheic and non-diarrheic calves. The veterinarian can use this as a starting point for further study in the future. The three pathogenic strains of *E. coli* that are significant for zoonotic diseases were identified in this study. Both diarrheic and non-diarrheic calves’ *E. coli* isolates that were resistant to different antibiotics were identified. This could serve as a warning against the improper use of antibiotics in the study area and used to enhance the economy of the country.

The study’s limitations include the inability to perform molecular detection on all *E. coli* isolates and the absence of virulence gene for all pathogenic *E. coli* strains during the study period. These impacted to identify the most common strains in the study area. The sample size for this study is small because it was done through purposive sampling. This could make it difficult to identify the factors that are genuinely connected to our result.

## 5. Conclusion and Recommendations

The results of this study showed that the prevalence of *E. coli* isolates in diarrheic calves was higher than in non-diarrheic calves. The season, age, house hygiene, health status, and first colostrum feeding time all influence the risk of *E. coli* infection in calves. Only one strain, STEC, was isolated from non-diarrheic calves; instances of STEC, EPEC, and EHEC were found in diarrheic calves. In the current study, high rate of multidrug-resistant was found in both healthy and diarrheic calves. Gentamicin, norfloxacin, and nitrofurantoin were the most effective antimicrobials in the study area. Thus, in light of the conclusion as mentioned earlier, the following recommendations are proposed:

➢ Farmers should be trained on the good calf management, quality and quantity of colostrum soon after birth and calf house and feeding equipment hygiene.
➢ Researchers should conduct more research into the prevalence of *E. coli* and the identification of pathogenic strains linked to diseases in order to develop control and prevention strategies.
➢ It is vital that researchers monitor the emergence and spread of antibiotic-resistant pathogens and use antibiotics appropriately.
➢ Government rules and regulations regarding the rational use of veterinary drugs should be followed by all relevant bodies.
➢ Further studies in vaccine design for *E. coli* should focus on the complexity and variability of its strains’ antigens and immune responses.
➢ Modern diagnostic tools are necessary to accurately identify pathogenic *E. coli*.

## 6. Acknowledgements

First and foremost, I would like to convey my sincere gratitude and respect to my admired main advisor, Prof. Endrias Zewdu, for his skilled assistance, strong interest, and unwavering support in my research activities. I just feel very lucky and fortunate to have had the opportunity to work under his excellent follow-up and direction. My appreciation also extends to my co-advisor, Dr. Kebede Abdisa, for his willingness and humility in all aspects, which included expert support with all research activities, editing my paper, and adding more pertinent details to strengthen and enhance the thesis. Also, I am very grateful to Mrs. Buzealem Asefa, who provided vital technical assistance during my conventional bacteriological activities at the Ambo University Zoonosis and Food Safety Laboratory.

More significantly, I want to thank the Department of Veterinary Laboratory Technology of Ambo University Mamo Mezemir Campus School of Veterinary Medicine for their kind assistance with reagents, equipment, and technical support during my study time and laboratory work. Deserving of special thanks is Dr. Wakuma Mitiku, the head of the Veterinary Laboratory Technology Department, for his devoted eagerness to help me in any way he can and for his constant good wishes for me.

My sincere gratitude goes out to the National Veterinary Institute (NVI) for allowing me to complete my molecular laboratory there. Additionally, I am also deeply indebted to Mr. Getaw Deresse for his patience and dedication in providing invaluable technical assistance throughout my stay in the molecular laboratory.

## 8 Annexes

### Annex 1: Questionnaire survey format

Data collection questionnaire regarding farm management systems related to the isolation of *E. coli* from calves in West Showa Zone, Ambo, Bako Tibe, and Ejere Woredas.

1. **Farm name Owners’ name (optional): _**

1.1. Address: Zone_Woreda_Kebele_When established_
2. **Overview of the farm**

2.1. Herd size: Cow_Calves_Heifer_Bulls
2.2. Calf sex: a) Male b) Female
2.3. Calf breed: a) Local b) Cross c) Exotic
2.4. Calf age group: a) 1-30days b) 31 -60days c) 61-90days d) 91days-120days
2.5. Production systems: a) Urban b) Pre-urban
3. **Farm management:**

3.1. Owner’s level of education: a) Uneducated b) Elementary school c) High school d) college graduate e) Animal production expert
3.2. Calf attendants): a) owner (household members) b) Other (by hired)
3.3. Sex of calf attendants: a) Male b) Female
3.4. Attendant level of education: a) a) Uneducated b) Elementary school c) High school d) Collage graduate d) Animal production expert
3.5. Pregnant cow caring: a) access to adequate nutrition and water b) Comfortable environment c) delivery pen d) the same barn
4. **Watering and feeding calves**

4.1. Knowledge on the value of colostrum for newborn calves: a) yes b) no
4.2. First colostrum feeding time: a) before 3hrs b) after 3hrs
4.3. Colostrum feeding method: a) suckling b) hand fed
4.4. Types of additional feed for the calves: a) Grazing b) Concentrates
4.5. Watering: a) free access b) providing by attendants (owners) periodically.
4.6. Source of water: a) tap water b) river c) open well d) other
4.7. Age of calf weaning: a) 2 weeks b) 3 weeks c) 4 weeks and above
5. **Calf housing**

5.1. Type of House: a) sheltered b) open-air
5.2. Calf pen location: a) within the cow house b) in an independent pen
5.3. Calf house bedding type: a) wood shaving b) straw bedding
5.4. Calf house cleaning: a) twice a day b) every day
5.5. Cleaning of cow teat before suckling/milking: a) yes b) no
5.6. Cleaning of calf feed and water equipment: a) yes b) no
6. **Calf health status**

6.1. Is calf diarrhea a farm issue? a) Yes b) no
6.2. Which age group has more diarrhea? a) Below 30days b) 31days–60days c) 61days–90days
6.3. Category of diarrhea: a) Yellowish b) Mucous c) Watery d) Bloody
7. **Knowledge of the owners for the antimicrobial and Treatment practice**

7.1. Antibiotic treatment for dry cow: a) yes b) no
7.2. Antibiotic therapy for diarrheic calf: a) Yes b) No
7.3. Is antibiotic used before diarrhea (as a prophylaxis)? a) Yes b) No
7.4. Who give antibiotics to the animals? a) Veterinarian b) non-veterinarian
7.5. Duration of antibiotic therapy: a) for one days b) three days c) five days
7.6. Drugs supply: a) Veterinary pharmacy, b) Local market
7.7. Frequently used antibiotics: ; ;
8. **Knowledge on the calf health, disease prevention, and control management**

8.1. What are significant health issues for the farm? --------------------------------
8.2. Is diarrhea a significant health issue for calves? -------------------------------
8.3. How disease is being prevented and controlled in the farm ------------------
8.4. Actions taken to isolate and treat sick calves -------------------------
8.5. How sick calves respond to treatment -------------------------------------

**Assessor**: Lalisa Chali

**Supervisors:** Professor Endirias Zewdu and Dr. Kebede Abdisa

### Annex 2: Fecal Sample Collecting Format

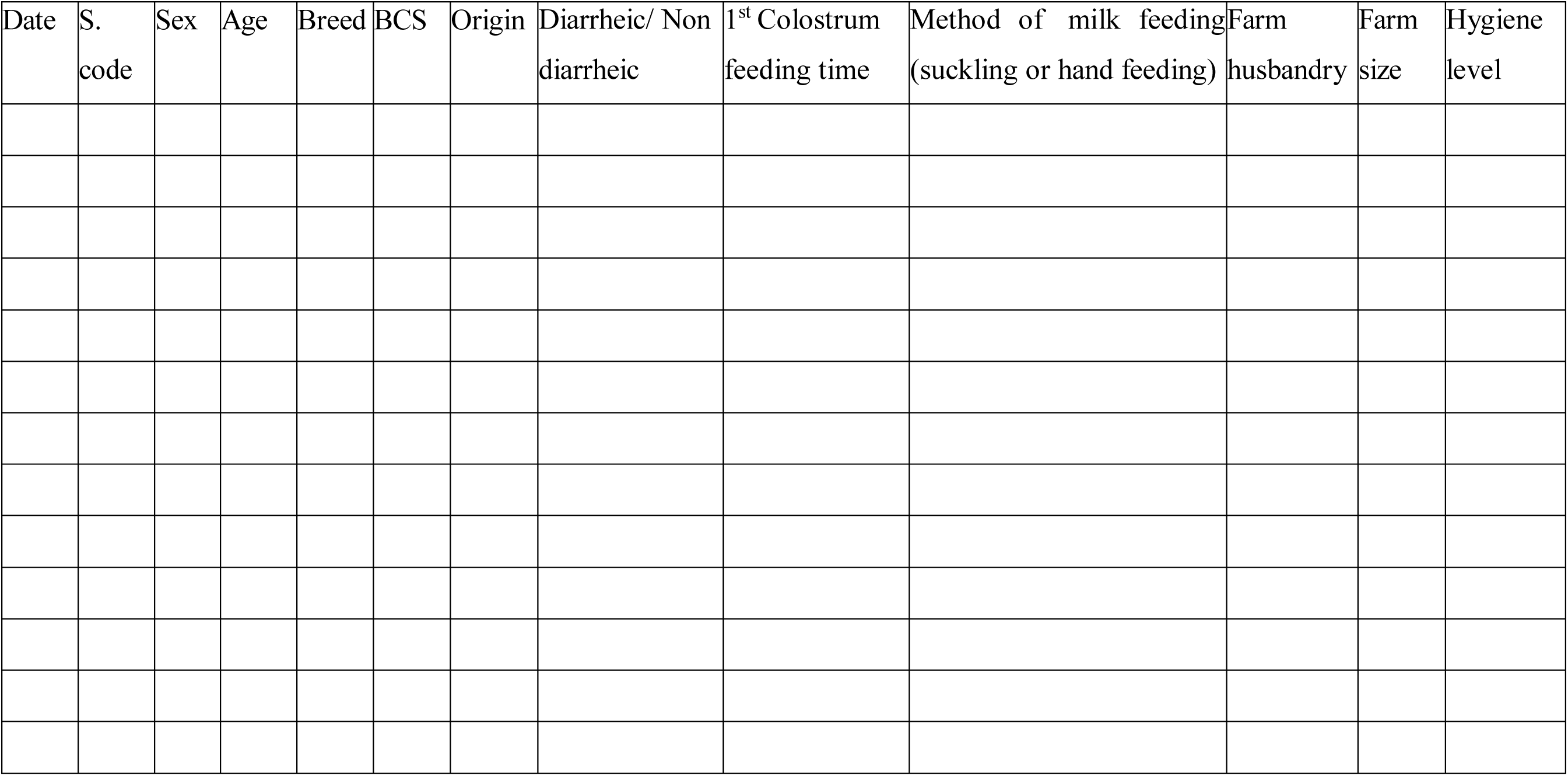

### Annex 3: Laboratory Recording Format

| Sample Code | Age | Sex | Diarrheic/non-diarrheic | MacConkey Pink/pale colony | EMB: Green metallic sheen/non-green metallic sheen colony | TSI (+/-) | IMViC Tests |  |  |  | Result (+/-) |
| --- | --- | --- | --- | --- | --- | --- | --- | --- | --- | --- | --- |
|  |  |  |  |  |  |  | Indole (+/-) | MR (+/-) | VP (+/-) | Citrate (+/-) |  |

### Annex 4: Lists and Preparation of Media

#### 1. Buffered peptone water (BPW) (HIMEDIA, India)

Compositions (g/liter): tryptones =10 g, sodium chloride = 5 g, disodium hydrogen phosephates. 12H2O = 9 g, potassium dihydrogen phosphate = 1.5 g, and water = 1000 ml. Preparation: 20.07 g of BPW is dissolved in 1000 ml of distilled water, mixed thoroughly, and sterilized by autoclaving at 121 °C for 15 minutes after being distributed in the test tubes.

#### 2. MacConkey Agar (HIMEDIA, India)

Compositions (g/liter): peptone = 17 g, proteose peptone = 3 g, lactose = 10 g, bile salts = 1.5 g, sodium chloride = 5 g, neutral red = 0.03 g, and agar = 15 g.

Preparation: 51.53 g is dissolved in 1000 ml of distilled water. Heat to boiling to dissolve the medium completely. It is sterilized by autoclaving at 15 psi of pressure with 121 °C for 15 minutes. Cool to 45 °C to 50 °C and pour into a sterile petri plate.

#### 3. Eosin Methylene Blue (EMB) Agar (HIMEDIA, Levine)

Compositions (g/liter): peptone = 10g, dipotassium phosphate = 2, lactose = 10g, eosin Y = 0.40g, methylene blue = 0065g, and agar = 15g

Preparation: 37.46 g of EMB dissolved in 1000 ml of distilled water. Sterilized by autoclaving at 15 psi pressure with 121°C for 15min. Cool to 45°C to 50°C and pour into a sterile petri dish. Precautions: Store the medium away from light to avoid photooxidation.

#### 4. Nutrient Agar (Accumix 500g, Spain)

Compositions: peptone = 5g, NaCl = 5g, beef extract = 1.5g, yeast extract = 1.5g, and agar = 15g.

Preparations: Suspend 28.0 g of nutrient agar powder in 1000 ml of distilled water. Mix thoroughly and boil with frequent agitation to dissolve the powder completely. Sterilized by autoclaving at 121 °C (15 psi) for 15 minutes.

#### 5. Triple sugar iron (TSI) agar (HIMEDIA, India)

Compositions (gms/liter): peptone = 10g, casein enzymatic hydrolysate = 10g, yeast extract = 3g, meat extract B = 3g, lactose = 10g, sucrose = 10g, dextrose = 1g, sodium chloride = 5g, ferrous sulfate = 0.20, sodium thiosulfate = 0.30g, phenol red = 0.024g, and agar = 12g.Preparations: Suspend 64.52 grams in 1000 ml of distilled water. Boil to dissolve the medium completely and distribute it into test tubes.

Sterilized by autoclaving at 15 psi pressure (121°C) for 15 minutes. Allow the medium to set in sloped form with a butt about 1 inch long.

#### 6. Indole test: Kovacs reagents (LOBA CHEMIA PVT. LTD., India)

Compositions: 1-butanol = 75 ml; dimethylaminobenzaldehyde = 0.92 g; HCl = 6.8 ml. Preparations: Mix the reagents with continuous stirring. 2 to 5 pure colonies were inoculated using a sterile wire loop in 2 ml of buffered peptone water. Incubated at 37°C for 24 hours, 0.5 ml of Kovac’s reagent was poured and observed after 2 minutes so that the reagent came to the top, and then the test culture was compared with the control tubes. The formation of a red or pink-colored ring at the top is taken as positive, while pale color or absence of red or pink is a negative indole test.

#### 7. MR-VP Medium (HiMedia, India)

Compositions (g/liter): buffered peptone = 7g, dextrose = 5g, dipotassium phosphate = 5g. Preparation: 17 g of this medium can be suspended by adding 1000 ml of distilled water, stirring well, and autoclaving for 15 minutes at 121°C.

#### 8. Simon Citrate Agar (Pronadisa, Spain)

Compositions (g/liter): Min assay = 99.5%, Chloride (Cl) = 0.001%, Sulfate (SO4) = 0.005%, Lead (Pb) = 0.005%, Calcium (Ca) = 0.002%, Iron (Fe) = 0.002%, Copper (Cu) = 0.001%, Sodium (Na) = 0.002%, Loss on drying (105°C) = 0.005%.

Preparations: Suspend 24.3 g of this medium in 1 liter of distilled water. Mix well and dissolve by heating with frequent agitation. Boil for 1 minute until complete dissolution. Dispense into the tube and sterilize in an autoclave at 121°C for 15 minutes. Allow it to cool in the standard position in order to obtain short butts of 1–1.5 cm in depth.

#### 9. Muller-Hinton Agar (Sisco Research Laboratories PVT. Ltd, India)

Compositions(g/l): beef infusion=17g; acid hydrolysate casein=17g; agar=17g; starch = 1.5g. Preparations: Add 39.0 g of powder to 1 liter of distilled or deionized water and mix carefully. Gently heat and bring to a boil. Autoclave at 15 psi pressure at 121 °C for 15 minutes.

### Annex 5: Protocols for Biochemical Tests

#### 1. Indole test

Principle: The bacteria that possess the tryptophanase enzyme can produce indole from tryptophane. When Kovac’s reagent is added to indole, it reacts to produce a red dye known as indole positive (+).

##### Procedures

- Isolated *E. coli* is inoculated in 2 ml of peptone water in a test tube.
- It should be incubated overnight at 37 ^°C.^
- 1 ml of Kovac’s reagent is added, and shake gently.
- Allow the tubes to stand for 2 minutes.
- Observe the reagent that comes to the top, forming a pink-red ring at the top.
- ***Escherichia coli*** is indole-positive.

#### 2. Methyl red (MR) Test

##### Principle

This is to detect the ability of an organism to produce and maintain stable acid end products from glucose fermentation. Some bacteria produce large amounts of acids from glucose fermentation that overcome the buffering action of the system. Methyl Red is a pH indicator, which remains red at a pH of 4.4 or less.

##### Procedures

- Isolated *E. coli* is inoculated into the MR-VP broth prepared.
- Incubated at 37 °C for 48 hours.
- The pH indicator of MR was added.
- The development of a red color is considered positive.
- MR-negative organisms produce a yellow color.
- ***Escherichia coli***: Positive for MR test

#### 3. Voges Proskauer (VP) test

##### Principle

It is employed to ascertain whether an organism ferments glucose to produce acetylmethyl carbinol. In the presence of atmospheric oxygen, ∝-naphthol, and a strong alkali (such as 40% KOH), acetylmethyl carbinol is converted to diacetyl. Naphthol acts as a color intensifier and must be added first. The compounds in the broth’s peptones that contain quinidine and diacetyl then condense to produce a pinkish-red color.

#### Procedures

- Isolated *E. coli* was inoculated into the MR-VP broth prepared.
- Incubated at 37°C for at least 48 hours.
- 6 ml of alpha-naphthol was added to the test broth and shaken.
- 2 ml of 40% KOH was added to the broth and shaken.
- The tube is allowed to stand for 15 minutes.
- The appearance of a red color was taken as a positive test.
- The negative tube color was yellow.
- ***Escherichia coli***: negative for the VP test

#### 4. Citrate utilization test

##### Principle

It detects an organism’s ability to utilize citrate as the sole source of carbon and energy. Procedures:

- *E. coli* colonies were inoculated into the slope of Simmon’s citrate agar.
- Incubated overnight at 37°C.
- If the organism can utilize citrate, the medium changes its color from green to blue.
- Observation: If the color of the medium changes to blue, it is citrate positive.
- ***Escherichia coli*:** Citrate negative

#### 5. Triple Sugar Iron (TSI) Tests

##### Principle

Ascertain the bacteria’s capacity to utilize lactose, sucrose, and glucose and how they interact with sodium thiosulfate. This makes it easier to distinguish between between differentiating bacterial isolates according to their capacity for fermentation and gas formation.

##### Procedures

- Isolated *E. coli* colony was inoculated by a wire loop piercing the agar butt at 3 to 5 mm depth.
- Gently streak the slant as you remove the wire to make sure the organism is exposed to aerobic conditions.
- Incubated at 37°C for at least 24 hours.
- See the medium for color variations on its slant and butt.
- Yellow slant and yellow butt (Acidic/Acidic) and gas formation (cracking medium)
- ***Escherichia coli***: Slant yellow, butt yellow (Y/Y) and gas production (cracks in agar)

### Annex 6: Disk Diffusion Test Procedures

#### Principles

It serves as a direction when selecting antibiotics to treat bacterial and fungal infections.

- Procedures: Four to five isolated colonies from pure fresh culture were transferred into tubes containing 5 ml of nutrient broth.
- The broth culture will be incubated at 37 °C for 4 to 6 hours until it achieves the 0.5 McFarland turbidity standards.
- A sterile cotton swab was dipped into the suspension, rotated several times, pressing firmly on the inside wall of the tube above the fluid level to remove excess inoculums, and swabbed uniformly over the surface of the Muller-Hinton agar plate.
- The plates were held at room temperature for 30 minutes to allow drying.
- Antimicrobial discs were placed on the inoculated agar surface by sterile forceps at about 2 cm apart and were placed at least 15 mm apart from the edge of the plates to prevent overlapping of the inhibition zones.
- The plates were incubated at 37 °C for 24 h aerobically.
- The diameter of the zones of inhibitions was recorded and classified as resistant, intermediate, or susceptible (CLSI, 2020).

### Annex 7: Recording Format for Antibiotic Susceptibility Test Results

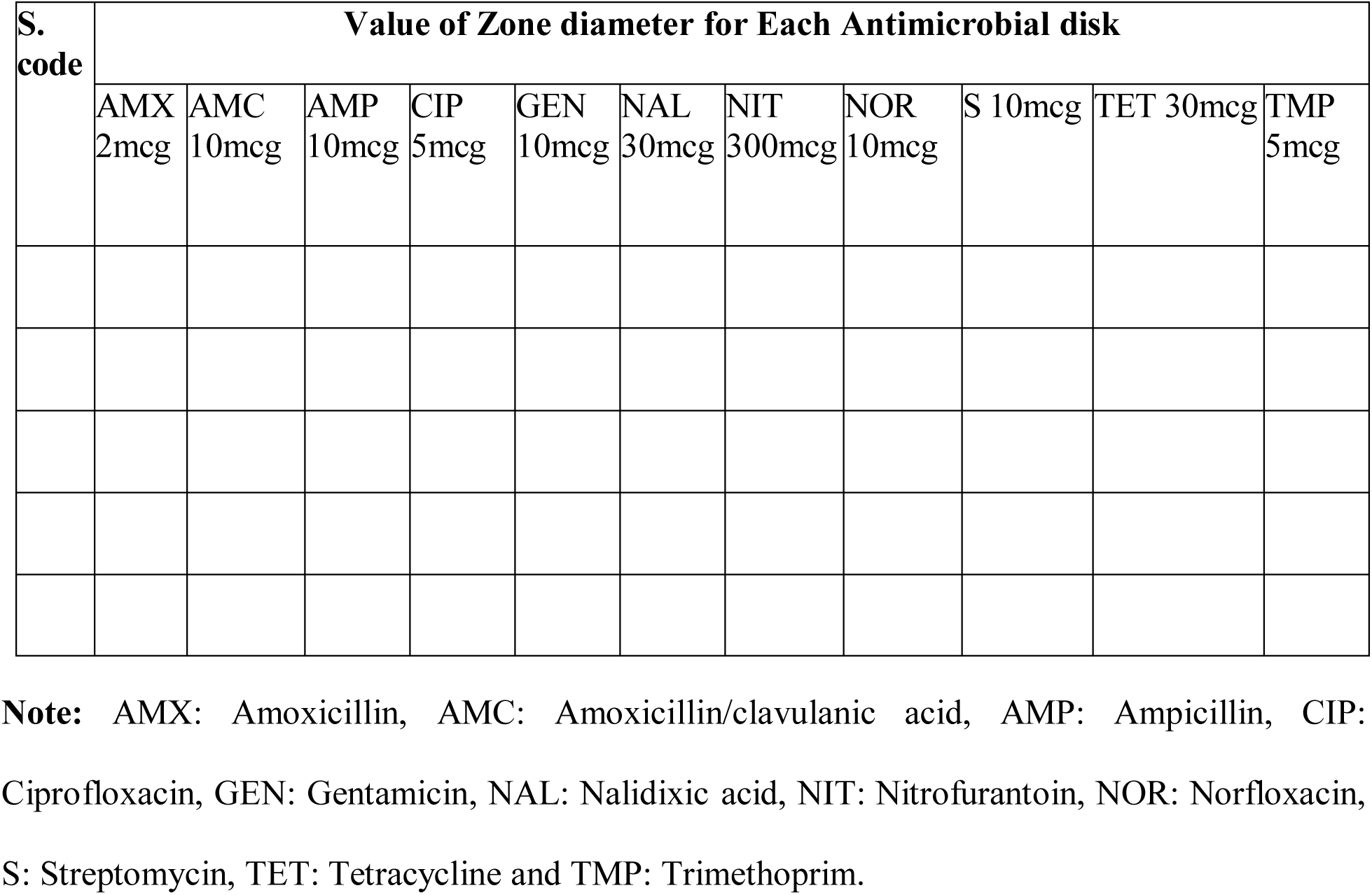

### Annex 8: Performance Standards for AMR Testing of *E. coli*

| No. | Antimicrobials | Susceptibility | Intermediate | Resistance |
| --- | --- | --- | --- | --- |
| 1 | Ciprofloxacin 5mg | $\geq 26$ | 22 – 25 | $\leq 21$ |
| 2 | Gentamicin 10mg | $\geq 15$ | 13 – 14 | $\leq 12$ |
| 3 | Nitrofurantoin 300mg | $\geq 17$ | 15 – 16 | $\leq 14$ |
| 4 | Norfloxacin 10mg | $\geq 17$ | 13 – 16 | $\leq 12$ |
| 5 | Nalidixic acid 30mg | $\geq 19$ | 14 – 18 | $\leq 13$ |
| 6 | Streptomycin 10mg | $\geq 15$ | 12 – 14 | $\leq 11$ |
| 7 | Amoxicillin/clavulanic acid 10mg | $\geq 18$ | 14 – 17 | $\leq 13$ |
| 8 | Amoxicillin 2mg | $\geq 17$ | 14 – 16 | $\leq 13$ |
| 9 | Ampicillin 10mg | $\geq 17$ | 12 – 16 | $\leq 13$ |
| 10 | Tetracycline 30mg | $\geq 15$ | 12 – 14 | $\leq 11$ |
| 11 | Trimethoprim 5mg | $\geq 16$ | 11 – 15 | $\leq 10$ |

Source: Clinical Laboratory Institute Standards (CLIS, 2020)

## Annex 9: Summary of the Overall Study

| No. | Activities | Description of Activities |
| --- | --- | --- |
| 1 | Fecal Sample collection | From diarrheic and non-diarrheic calves |
| 2 | Pre-enrichment | 25g of fecal sample in 225ml Buffered Peptone Water or at 1:9 ratios incubated at 37°C for 24hrs. |
| 3 | Culturing on selective media | A loop full of enriched culture broths on MacConkey agar with an inoculation loop incubated at 37°C for 24-48hrs |
|  |  | Assumed isolates from MacConkey agar to Eosin methylene blue (EMB) plate cultured in an inoculation loop at 37°C for 24 to 48 hours. |
|  |  | Isolated colonies were kept on nutrient agar for a full day at 37 °C and used for biochemical tests. |
| 4 | Biochemical tests |  |
| 4.1 | Triple sugar iron agar test | Pick an isolated colony from Nutrient agar and inoculate it with a straight inoculating wire in the middle of the agar. Incubated at 37 °C for 18 hours. |
| 4.2 | IMVIC test |  |
| 4.2.1 | Indole test | Isolated *E. coli* was inoculated into peptone water. And incubated at 37 °C for 48 hours. |
| 4.2.2 | Methyl red (MR) test | Isolated *E. coli* was inoculated in MR-VP broth. And incubated at 37 °C for 48 hours. |
| 4.2.3 | Voges Proskauer (VP) test | Isolated *E. coli* was inoculated in MR-VP broth. And incubated at 37 °C for 48 hours. |
| 4.2.4 | Citrate utilization test | Isolated *E. coli* was inoculated in a citrate slant. And incubated at 37 °C for 48 hours. |
| 5 | Virulent Gene Detection | DNA extracted using the boiling method |
|  |  | DNA amplification by PCR |
|  |  | Analyzed by gel electrophoresis |
| 6 | Antimicrobial sensitivity test | Fresh cultures were transferred into a test tube of 5 ml of Nutrient broth was incubated at 37 °C for 6 hours. |
|  |  | The culture was wiped onto Mueller-Hinton agar plates, an antibiotic disk was added. The plates were incubated for twenty-four hours at 37 °C. |

## Annex 10: A List of Pictures taken throughout the Study

**Figures 1:**
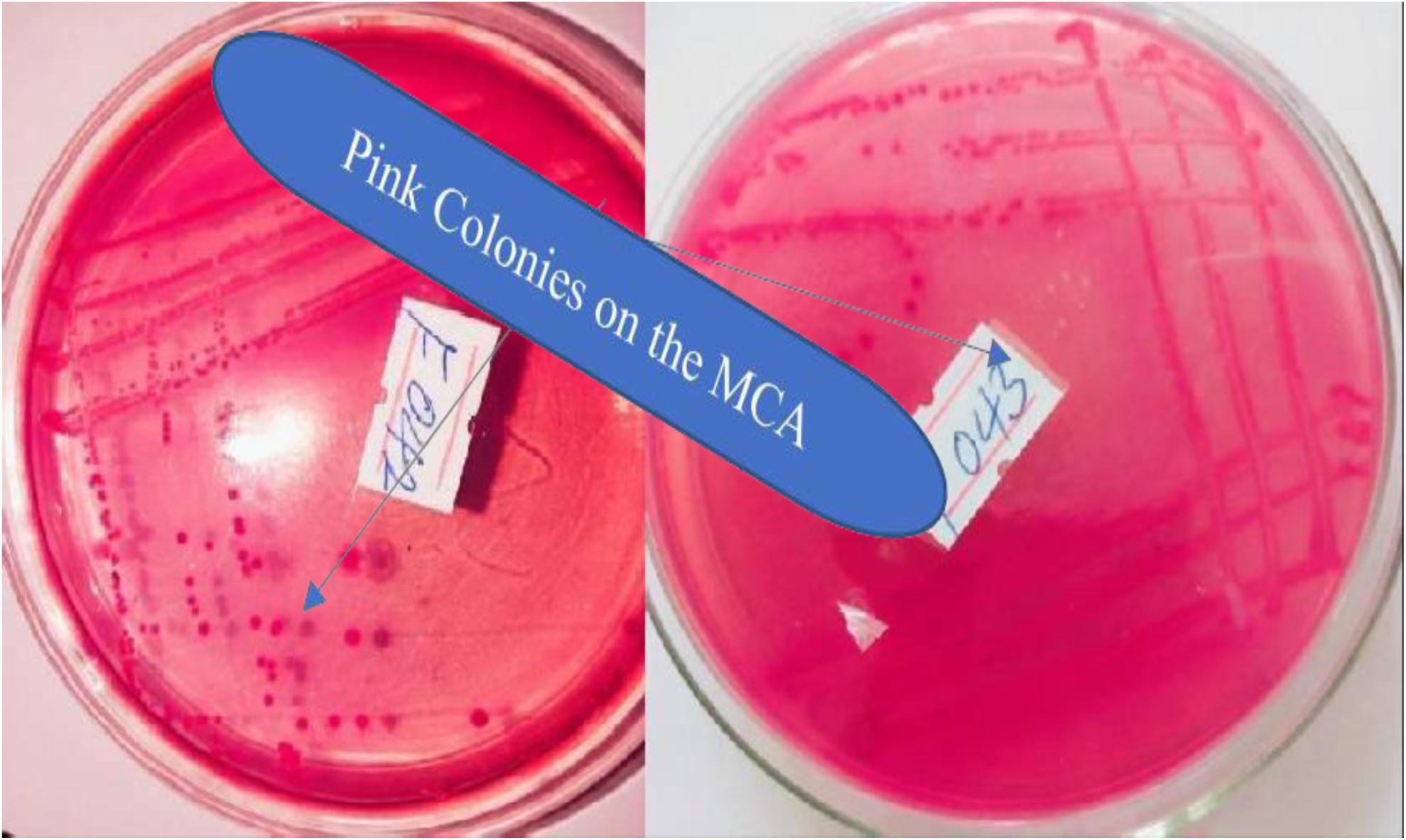
Colony growth appearance on MacConkey agar.

**Figures 2:**
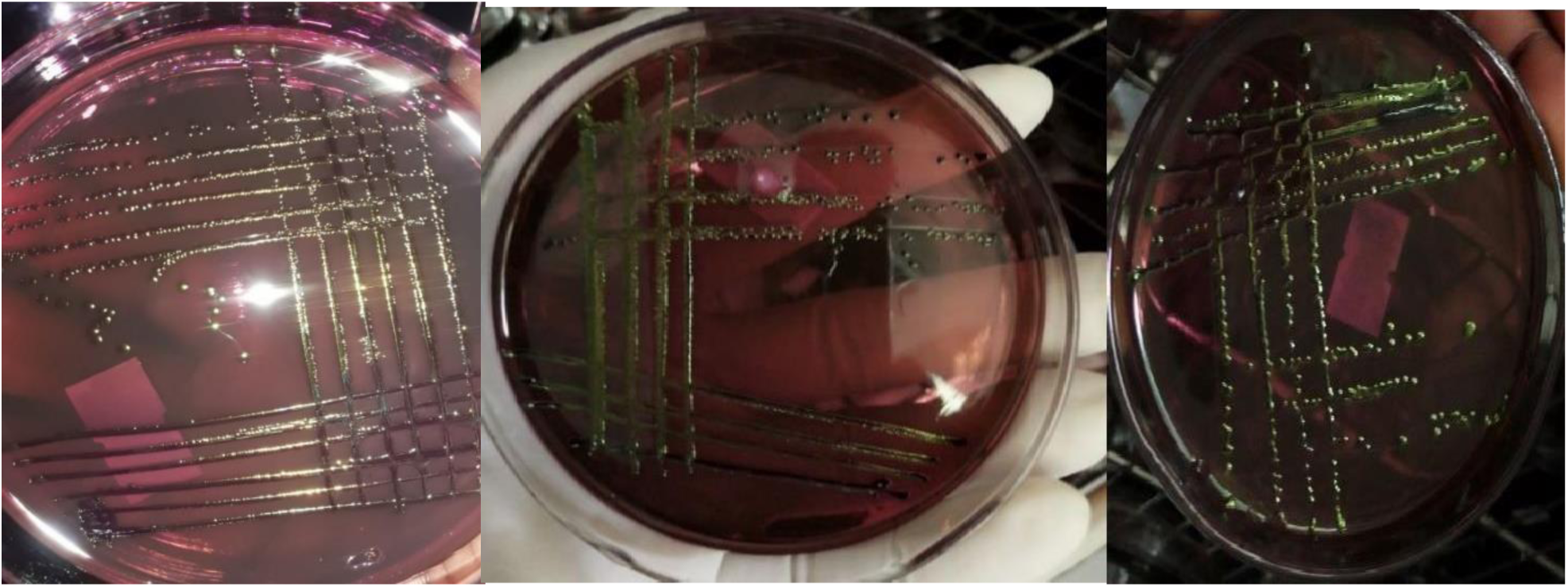
Growth appearance of the colonies on the EMB Plate.

**Figures 3:**
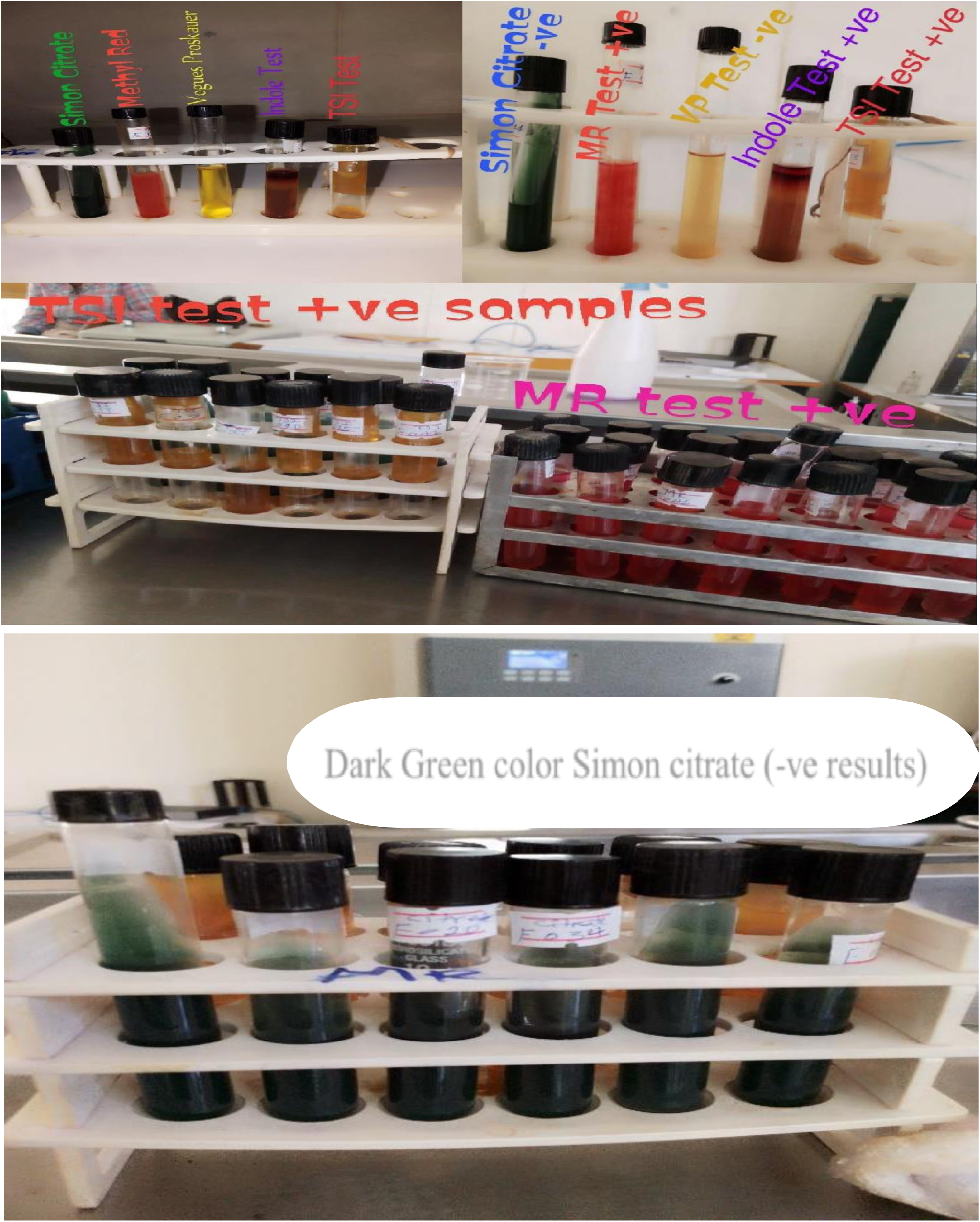
Biochemical Tests (TSI and IMViC)

## References

1. Merera O, Regassa A, Gizaw F, Adugna B, Abera S. Isolation, Identification and Antimicrobial Susceptibility Tests of *Escherichia coli* and *Salmonella* from diarrheic calves in and around Sebeta Town 1–18. 2020.

2. Tamiru M, Amza N. Review on the status of dairy cattle production in Ethiopia. Journal of Genetic and Environmental Resources Conservation. 2017;5(2):84–95.

3. Fentie T, Guta S, Mekonen G, Temesgen W, Melaku A, Asefa G, Tesfaye S, Niguse A, Abera B, Kflewahd FZ, Hailu B. Assessment of major causes of calf mortality in urban and periurban dairy production system of Ethiopia. Veterinary Medicine International. 2020;2020(1):3075429.

4. Khawaskar DP, Sinha DK, Lalrinzuala MV, Athira V, Kumar M, Chhakchhuak L, Mohanapriya K, Sophia I, Abhishek, Kumar OV, Chaudhuri P. Pathotyping and antimicrobial susceptibility testing of *Escherichia coli* isolates from calves. Veterinary Research Communications. 2022 Jun;46(2):353–62.

5. Ramos S, Silva V, Dapkevicius MD, Caniça M, Tejedor-Junco MT, Igrejas G, Poeta P. *Escherichia coli* as commensal and pathogenic bacteria among food-producing animals: Health implications of extended spectrum β-lactamase (ESBL) production. Animals. 2020 Nov 29;10(12):2239.

6. Chekole WS, Adamu H, Sternberg-Lewrein S, Magnusson U, Tessema TS. Occurrence of *Escherichia coli* pathotypes in diarrheic calves in a low-income setting. Pathogens. 2022 Dec 27;12(1):42.

7. Tedla M, Degefa K. Bacteriological study of calf coli septicemia in Alage dairy farm, southern Ethiopia. BMC research notes. 2017 Dec; 10:1–5.

8. Muktar YI, Bishoftu E. Major Enteropathogens Associated in Calf Diarrhoea with an Emphasis on *Escherichia coli* and *Salmonella* Species in Dairy Farms of Muke Turi, Debre Stige and Fitche towns North Shewa, Ethiopia.

9. Croxen MA, Law RJ, Scholz R, Keeney KM, Wlodarska M, Finlay BB. Recent advances in understanding enteric pathogenic *Escherichia coli*. Clinical microbiology reviews. 2013 Oct;26(4):822–80.

10. Conrad CC, Stanford K, Narvaez-Bravo C, Callaway T, McAllister T. Farm fairs and petting zoos: a review of animal contact as a source of zoonotic enteric disease. Foodborne pathogens and disease. 2017 Feb 1;14(2):59–73.

11. Elmonir W, Abo-Remela E, Sobeih A. Public health risks of *Escherichia coli* and Staphylococcus aureus in raw bovine milk sold in informal markets in Egypt. The Journal of Infection in Developing Countries. 2018 Jul 31;12(07):533–41.

12. Gambushe SM, Zishiri OT, El Zowalaty ME. Review of *Escherichia coli* O157: H7 prevalence, pathogenicity, heavy metal and antimicrobial resistance, African perspective. Infection and Drug Resistance. 2022 Jan 1:4645–73.

13. Mohammed SA, Marouf SA, Erfana AM, El JK, Hessain AM, Dawoud TM, Kabli SA, Moussa IM. Risk factors associated with *Escherichia coli* causing neonatal calf diarrhea. Saudi journal of biological sciences. 2019 Jul 1;26(5):1084–8.

14. Gebregiorgis A, Tessema TS. Characterization of *Escherichia coli* isolated from calf diarrhea in and around Kombolcha, South Wollo, Amhara Region, Ethiopia. Tropical animal health and production. 2016 Feb; 48:273–81.

15. Duse A, Waller KP, Emanuelson U, Unnerstad HE, Persson Y, Bengtsson B. Risk factors for antimicrobial resistance in fecal *Escherichia coli* from pre-weaned dairy calves. Journal of dairy science. 2015 Jan 1;98(1):500–16.

16. Cao H, Bougouffa S, Park TJ, Lau A, Tong MK, Chow KH, Ho PL. Sharing of AMR genes between humans and food animals. M-Systems. 2022 Dec 20;7(6): e00775–22.

17. Muktar Y, Mamo G, Tesfaye B, Belina D. A review on major bacterial causes of calf diarrhea and its diagnostic method. Journal of Veterinary Medicine and Animal Health. 2015 May 31;7(5):173–85.

18. Tarekegn Y, Molla F. The Prevalence of *Escherichia coli* from diarrheic calves and their antibiotic sensitivity test in selected dairy farms of Debre Zeit, Ethiopia. Adv Biotechnol Microbiol. 2017;6(1):555–9.

19. Wale Y, Kassa T. Antimicrobial susceptibility pattern of *Escherichia coli* isolated from dairy calves with diarrhoea in Akaki Kality, Addis Ababa, Ethiopia. Journal of Applied Animal Research. 2023 Dec 31;51(1):470–6.

20. Akane AE, Tessema TS, Ali DA. Distribution of *Escherichia coli* biotypes shed by dairy calves in selected dairy farms in Bishoftu, Ethiopia. Ethiopian Veterinary Journal. 2022 Aug 29;26(2):132–42.

21. Loayza F, Graham JP, Trueba G. Factors obscuring the role of *Escherichia coli* from domestic animals in the global antimicrobial resistance crisis: an evidence-based review. International journal of environmental research and public health. 2020 May;17(9):3061.

22. Gemeda BA, Assefa A, Jaleta MB, Amenu K, Wieland B. Antimicrobial resistance in Ethiopia: A systematic review and meta-analysis of prevalence in foods, food handlers, animals, and the environment. One Health. 2021 Dec 1; 13:100286.

23. Belete MA, Demlie TB, Chekole WS, Sisay Tessema T. Molecular identification of diarrheagenic *Escherichia coli* pathotypes and their antibiotic resistance patterns among diarrheic children and in contact calves in Bahir Dar city, Northwest Ethiopia. PLOS One. 2022 Sep 28;17(9): e0275229.

24. ECDC. Antimicrobial Resistance in the EU/EEA (EARS-Net) Annual Epidemiological Report for 2019. Epidemiology of antimicrobial resistance EU/EEA. 2020; 174:341.

25. Zhou Y, Li Y, Zhang L, Wu Z, Huang Y, Yan H, Zhong J, Wang LJ, Abdullah HM, Wang HH. Antibiotic administration routes and oral exposure to antibiotic resistant bacteria as key drivers for gut microbiota disruption and resistome in poultry. Frontiers in Microbiology. 2020 Jul 7; 11:1319.

26. Mesele F, Leta S, Amenu K, Abunna F. Occurrence of *Escherichia coli* O157: H7 in lactating cows and dairy farm environment and the antimicrobial susceptibility pattern at Adami Tulu Jido Kombolcha District, Ethiopia. BMC Veterinary Research. 2023 Jan 11;19(1):6.

27. Endale A. Characterization of Diarrheagenic *Escherichia coli* strains isolated from dairy calves in Fiche, Debretsige and Muketuri towns. North Shoa: MSc thesis, Addis Ababa University; 2018. Available from: http://etd.aau.edu.et/handle/123456789/18210

28. Adisu R, Kebed A. Observational Study of Major Dairy Health Problems in Ambo and Holeta Town, Oromia Region. East Afri. Scholars J. Agri. Life Sci. 2018;1(1):33–8.

29. Abebaye H, Mengistu A, Tamir B, Assefa G, Feyisa F. Feed resources availability and feeding practices of smallholder farmers in selected districts of West Shewa Zone, Ethiopia. World J Agric Sci. 2019;15(1):21–30.

30. Girma A, Workineh C, Bekele A, Tefera D, Gelana H, Gebisa D, Chalchisa T, Pal M. Assessment of farmers knowledge and attitude on vaccination of livestock and its implications in Ejere district of West Shewa Zone, Oromia, Ethiopia. Acta Scientific microbiology (ISSN: 2581-3226). 2022 Dec;5(12).

31. Hordofa D, Abunna F, Megersa B, Abebe R. Incidence of morbidity and mortality in calves from birth to six months of age and associated risk factors on dairy farms in Hawassa city, southern Ethiopia. Heliyon. 2021 Dec 1;7(12).

32. Birhanu F, Lemma D, Eticha E, Abera B, Adem A. Prevalence and risk factors of cryptosporidiosis in calves in Asella town, south eastern, Ethiopia. Acta Parasitologica Globalis. 2017;8(1):50–7.

33. Coura FM, de Araújo Diniz S, Mussi JM, Silva MX, Lage AP, Heinemann MB. Characterization of virulence factors and phylogenetic group determination of *Escherichia coli* isolated from diarrheic and non-diarrheic calves from Brazil. Folia microbiologica. 2017 Mar; 62:139–44.

34. Zucali M, Bava L, Tamburini A, Guerci M, Sandrucci A. Management risk factors for calf mortality in intensive Italian dairy farms. Italian Journal of Animal Science. 2013 Jan 1;12(2): e26.

35. Messele YE, Hasoon MF, Trott DJ, Veltman T, McMeniman JP, Kidd SP, Low WY, Petrovski KR. Longitudinal analysis of antimicrobial resistance among Enterococcus species isolated from Australian beef cattle Faeces at feedlot entry and exit. Animals. 2022 Oct 6;12(19):2690.

36. Yeshiwas T, Fentahun WM. The prevalence of *Escherichia coli* from diarrheic calves and their antibiotic sensitivity test in selected dairy farms of Debre Zeit, Ethiopia. Adv Biotechnol Microbiol. 2017;6(1):555680. doi:10.19080/AIBM.2017.06.555680.

37. Abdlla YA, Al-Sanjary RA. The molecular identification of diarrheagenic *Escherichia coli* isolated from meat and meat products. Iraqi J Vet Sci. 2023;37(1):9–15.

38. Salama M, Younis W, Mohamed H, Sultan S. Microbiological and molecular characterization of *Escherichia coli* and *Salmonella* isolated from diarrheic calves. SVU-Int J Vet Sci. 2023;6(4):1–14.

39. Songsri J, Mala W, Wisessombat S, Siritham K, Cheha S, Noisa N, Klangbud WK. First isolation of verocytotoxin-producing *Escherichia coli* O157:H7 from sports animals in Southern Thailand. Vet World. 2022;15(9):2275.

40. Clinical and Laboratory Standards Institute (CLSI). Performance standards for antimicrobial susceptibility testing. 30th ed. CLSI document M100. Wayne (PA): CLSI; 2020. In: CLSI, 40(1).

41. Tadesse N. Prevalence and multidrug resistance profiles of *Escherichia coli* in dairy farms. Int J Vet Sci Res. 2020;6(2):142–147.

42. Ryu JH, Kim S, Park J, Choi KS. Characterization of virulence genes in *Escherichia coli* strains isolated from pre-weaned calves in the Republic of Korea. Acta Vet Scand. 2020;62(1):1–7.

43. El Bably MA, Mohammed AN, Mohamed MB, Fahmy HA. Monitoring and molecular characterization of multidrug resistant enteropathogenic *Escherichia coli* in dairy calves and their environment. J Vet Med Res. 2016;23(2):155–167.

44. Himani KM, Nayak A, Jogi J, Shakya P, Bordoloi S, Lade A, Gupta B. Isolation, identification, and antibiotic sensitivity pattern of *Escherichia coli* from diarrheic and non-diarrheic calves. Int J Curr Microbiol Appl Sci. 2021;10(01):2079–2085.

45. M ELHADY, A.H.M.E.D., M EL-AZZOUNY, M.O.N.A. and H ABOU-KHADRA, S.A.L.L.Y., 2020. Factors affecting calf enteritis infection caused by *Salmonellae* and *Escherichia coli*. Assiut Veterinary Medical Journal, 66(165), pp.21–43.

46. Bist B, Sharma B, Kumar A, Singh S, Jain U, Goswami M, Basak G. Seasonal effect on the prevalence of virulence genes of non-O157 verotoxigenic *Escherichia coli* serogroups in feces of cattle calves. Indian J Anim Sci. 2023;93(11):1046–1052.

47. Zaki MS, Abd-El-All AM, Attia AS, Dahshan H, Al-Ashery MA, Megahed A. *Escherichia coli* and Salmonella enterica isolated from Egyptian dairy cattle herds: the prevalence and molecular characteristics. Open Vet J. 2024;14(1):214.

48. Ali DA, Tesema TS, Belachew YD. Molecular detection of pathogenic *Escherichia coli* strains and their antibiogram associated with risk factors from diarrheic calves in Jimma Ethiopia. Sci Rep. 2021;11(1).

49. Umpiérrez A, Ernst D, Fernández M, Oliver M, Casaux ML, Caffarena RD, Zunino P. Virulence genes of *Escherichia coli* in diarrheic and healthy calves. Rev Argent Microbiol. 2021;53(1):34–38.

50. Fecteau G, Fairbrother JM, Higgins R, Van Metre DC, Paré J, Smith BP, Jang S. Virulence factors in *Escherichia coli* isolated from the blood of neonatal calves. Vet Microbiol. 2001;78(3):241–249.

51. Morin MP, Dubuc J, Freycon P, Buczinski S. A calf-level study on colostrum management practices associated with adequate transfer of passive immunity in Québec dairy herds. J Dairy Sci. 2021;104(4):4904–4913.

52. Lichtmannsperger K, Hartsleben C, Spöcker M, Hechenberger N, Tichy A, Wittek T. Factors associated with colostrum quality, the failure of transfer of passive immunity, and the impact on calf health in the first three weeks of life. Animals. 2023;13(11):1740.

53. Fischer AJ, Song Y, He Z, Haines DM, Guan LL, Steele MA. Effect of delaying colostrum feeding on passive transfer and bacterial colonization in neonatal male Holstein calves. J Dairy Sci. 2018;101(4):3099–3109.

54. Tutija JF, Ramos CA, Lemos RA, Santos AA, Reckziegel GH, Freitas MG, Leal CR. Molecular and phenotypic characterization of *Escherichia coli* from calves in an important meat-producing region in Brazil. J Infect Dis Control. 2022 Jun 30;16(6):1030-1036. Doi: 10.3855/jidc.13377.

55. Hosein HI, Azzam RA, Abo-Elwafa M, Menshawy AMS, Rouby S. Virulence profile of enteropathogenic *Escherichia coli* (EPEC) isolated from the cases of neonatal calf diarrhea. Adv Anim Vet Sci. 2019;7(9):755–760.

56. Kolenda R, Burdukiewicz M, Schierack P. A systematic review and meta-analysis of the epidemiology of pathogenic *Escherichia coli* of calves and the role of calves as reservoirs for human pathogenic *Escherichia coli*. Front Cell Infect Microbiol. 2015; 5:23.

57. Padola NL, Castro V, Etcheverría A, Figueiredo E, Guillén R, Umpiérrez A. Bovine reservoir of STEC and EPEC: advances and new contributions. In: Trending topics in Escherichia coli research: the Latin American perspective. Cham: Springer International Publishing; 2023. p. 107–127.

58. Keykhaei N, Salari S, Rashki A. Frequency of K99, stx1, and stx2 virulence factors in *Escherichia coli* isolated from diarrheic and clinically healthy suckling calves in Sistan and Baluchistan Province, Iran. Arch Razi Inst. 2021;76(2):283.

59. Gomes V, Barros BP, Castro-Tardon DI, Martin CC, Santos FCR, Knöbl T, Hurley DJ. The role of anti-*Escherichia coli* antibody from maternal colostrum on the colonization of newborn dairy calves’ gut with *Escherichia coli* and the development of clinical diarrhea. Animal (Basel). 2023; 2:100037.

60. Mathew X, Janus A, Deepa PM, Bipin KC, Habeeb BP, Vergis J. Molecular characterization of *E. coli* from colibacillosis affected calves, Wayanad, Kerala. 2020;9(8):141–144.

61. Costa LRM, Buiatte ABG, da Cunha Dias S, Garcia LNH, Cossi MVC, Yamatogi RS, Pereira JG. Influence of animal production systems on the presence of pathogenic strains of *Escherichia coli* in the bovine production chain. Food Control. 2024; 157:110155.

62. Madoshi BP, Kudirkiene E, Mtambo MM, Muhairwa AP, Lupindu AM, Olsen JE. Characterization of commensal *Escherichia coli* isolated from apparently healthy cattle and their attendants in Tanzania. PLoS One. 2016;11(12): e0168160.

63. Hang BPT, Wredle E, Börjesson S, Sjaunja KS, Dicksved J, Duse A. High level of multidrug-resistant *Escherichia coli* in young dairy calves in southern Vietnam. Trop Anim Health Prod. 2019; 51:1405–1411.

64. Geletu US, Usmael MA, Ibrahim AM. Isolation, identification, and susceptibility profile of *E. coli*, *Salmonella*, and *S. aureus* in dairy farm and their public health implication in Central Ethiopia. BioMed Res Int. 2022; 2022:1887977. Doi: 10.1155/2022/1887977.

65. Mulu BM, Belete MA, Demlie TB, Tassew H, Sisay Tessema T. Characteristics of pathogenic *Escherichia coli* associated with diarrhea in children under five years in northwestern Ethiopia. Trop Med Infect Dis. 2024;9(3):65.

66. Dulo F, Feleke A, Szonyi B, Fries R, Baumann MP, Grace D. Isolation of multidrug-resistant *Escherichia coli* O157 from goats in the Somali region of Ethiopia: a cross-sectional, abattoir-based study. PLoS One. 2015;10(11): e0142905.

67. Bedasa S, Shiferaw D, Abraha A, Moges T. Occurrence and antimicrobial susceptibility profile of *Escherichia coli* O157:H7 from food of animal origin in Bishoftu town, Central Ethiopia. Int J Food Contam. 2018;5(1):1–8.

68. Sarba EJ, Kelbesa KA, Bayu MD, Gebremedhin EZ, Borena BM, Teshale A. Identification and antimicrobial susceptibility profile of *Escherichia coli* isolated from backyard chicken in and around Ambo, Central Ethiopia. BMC Vet Res. 2019; 15:85.

69. Sarba EJ, Wirtu W, Gebremedhin EZ, Borena BM, Marami LM. Occurrence and antimicrobial susceptibility patterns of *E. coli* and *E. coli* O157 isolated from cow milk and milk products, Ethiopia. Scientific Reports. 2023;13(1):16018.

70. Tadesse T, Alemayehu H, Medhin G, Akalu A, Eguale T. Antibiogram of *Escherichia coli* isolated from dairy cattle and in-contact humans in selected areas of Central Ethiopia. Vet Med Res Rep. 2024;117–127.

71. El-shazly WSA, El-Tawab A, Awad A, Elhofy FI, El Hamalawy AA, Abo El Ella AA, El-khayat ME. Prevalence of multi drug resistant *Escherichia coli* in diarrheic ruminants. Benha Vet Med J. 2020;38(1):75–78.

72. Balcha FB, Sulayeman M, Neja SA. Prevalence and antimicrobial susceptibility of *Staphylococcus aureus* and *Escherichia coli* isolates of bovine mastitis and associated risk factors in Shashemene Town, Ethiopia. J Vet Heal Sci. 2022;3(4):361–372.

73. Fesseha H, Mathewos M, Aliye S, Mekonnen E. Isolation and antibiogram of *Escherichia coli* O157:H7 from diarrheic calves in urban and peri-urban dairy farms of Hawassa town. Vet Med Sci. 2022;8(2):864–876.

74. Arbab S, Ullah H, Wei X, Wang W, Ahmad SU, Zhang J. Drug resistance and susceptibility testing of Gram-negative bacterial isolates from healthy cattle with different β–Lactam resistance phenotypes from Shandong province China. Brazilian Journal of Biology. 2021;83: e247061.

75. Bag MAS, Khan MSR, Sami MDH, Begum F, Islam MS, Rahman MM, Hassan J. Virulence determinants and antimicrobial resistance of *E. coli* isolated from bovine clinical mastitis in some selected dairy farms of Bangladesh. Saudi J Biol Sci. 2021;28(11):6317–6323.

76. Hoyle DV, Yates CM, Chase-Topping ME, Turner EJ, Davies SE, Low JC, Amyes SG. Molecular epidemiology of antimicrobial-resistant commensal *Escherichia coli* strains in a cohort of newborn calves. Appl Environ Microbiol. 2005;71(11):6680–6688.

77. Chirila F, Tabaran A, Fit N, Nadas G, Mihaiu M, Tabaran F, Dan SD. Concerning the increase in antimicrobial resistance in Shiga toxin-producing *Escherichia coli* isolated from young animals during 1980–2016. Microbes Environ. 2017;32(3):252–259.

